# Twisting of the heart tube during cardiac looping is a *tbx5*-dependent and tissue-intrinsic process

**DOI:** 10.1101/2020.08.03.230359

**Authors:** Federico Tessadori, Fabian Kruse, Susanne C. van den Brink, Malou van den Boogaard, Vincent M. Christoffels, Jeroen Bakkers

## Abstract

Organ laterality refers to the Left-Right (LR) asymmetry in disposition and conformation of internal organs, established in the developing embryo. The heart is the first organ to display visible LR asymmetries as it is positioned to the left side of the midline and undergoes rightward looping morphogenesis. Cardiac looping morphogenesis is tightly controlled by a combination of heart-intrinsic and -extrinsic mechanisms. As the mechanisms that drive cardiac looping are not well understood, we performed a forward genetic screen for zebrafish mutants with defective heart looping. We describe a new loss-of-function allele for *tbx5a*, which displays normal leftward positioning but defective rightward looping morphogenesis. By using live two-photon confocal imaging to map cardiomyocyte behavior during cardiac looping at a single-cell level we establish that during looping, ventricular and atrial cardiomyocytes rearrange in opposite directions towards the outer curvatures of the chambers. As a consequence, the cardiac chambers twist around the atrioventricular canal resulting in torsion of the heart tube, which is compromised in *tbx5a* mutants. Manipulations of cardiac looping by chemical treatment and *ex vivo* culture establishes that the twisting of the heart tube depends on intrinsic mechanisms and is independent from tissue growth by cell addition. Furthermore, the cardiac looping defect in *tbx5a* mutants is rescued in *tbx5a/tbx2b* double mutants, indicating that it requires proper tissue patterning. Together, our results establish that cardiac looping in zebrafish involves twisting of the chambers around the AV canal, which requires correct tissue patterning by Tbx5a.

## Introduction

Bilateral animals such as vertebrates, while being symmetric on the outside when divided through the sagittal plane, have left-right (LR) asymmetrically arranged internal organs. LR asymmetry of organ disposition and form supports proper development and function of the organism throughout life.

The embryonic heart is the first organ to visibly break LR symmetry of the vertebrate embryo ((1) and references therein). The heart starts out as a linear tube, which is tilted towards the left relative to the midline in mouse and zebrafish. This is followed by an asymmetric bending towards the right, initiating an ensemble of developmentally-regulated complex processes referred to as cardiac looping (2). The looped heart tube is either a flat S-shape in fish or a helix in amniotes (chick and mouse) (1). Correct looping is closely intertwined to proper patterning and alignment of the inflow and outflow tracts, cardiac chambers and atrioventricular canal, which are crucial to establish and maintain heart function. Indeed, cardiac looping defects in humans can result in severe congenital heart defects such as transposition of the great arteries (TGA), double outlet right ventricle (DORV) and Tetralogy of Fallot (TOF)(3). Correct cardiac looping depends on both tissue intrinsic and extrinsic mechanisms. Establishment of LR asymmetry involves an extrinsic mechanism that influences cardiac looping. In most vertebrates this LR asymmetry is established during embryogenesis due to the activity of the LR organizer, called the node in mice and Kupffer’s vesicle in zebrafish. The LR organizer is a transient structure consisting of ciliated cells, located in the posterior part of the embryo (4). Rotation of the cilia results in a directed fluid flow (nodal flow), which breaks the symmetry by inducing left-sided specific expression of Nodal and Pitx2 (5, 6). Left-sided Nodal expression regulates the asymmetric position and dextral looping of the heart (5, 7–10). In zebrafish, LR symmetry is first broken when the linear heart tube arises from an initial flat disc between 20- and 24-hours post-fertilization (hpf; reviewed in (11)). As its formation progresses, the inflow pole moves to the left side of the midline in a process referred to as cardiac jogging (12). This breaking of LR asymmetry is dependent on left-sided Nodal expression (8, 13, 14). After this, the heart tube undergoes cardiac looping, which under normal conditions is dextral (rightward). If the function of the LR organ is affected, a sinistral (leftward) loop can be observed (9) (15). Based on mutant analysis it was suggested that cardiac jogging can be separated from cardiac looping and that there are likely separate mechanisms that regulate these processes (reviewed by (16)). Corroborating such as model, we previously demonstrated that while left-sided Nodal expression directs cardiac jogging, a separate, tissue-intrinsic mechanism drives looping morphogenesis (9). Intrinsic LR asymmetry has been observed in various tissues and organs of invertebrates (reviewed in (17)). In Drosophila the hindgut and the genitalia show LR asymmetry (18, 19), for which myosin seems to be the major determinant (20, 21). LR asymmetry is not only observed at the organ and tissue level, but also in single cells (reviewed in (22)). For example, human leukemia cells preferentially polarize to the left of an imaginary axis between the nucleus and the centrosome (23). The actin cytoskeleton and actomyosin interactions are important for the observed intrinsic chirality of cells (reviewed in (24)) as chiral actin cytoskeletal organization was observed in cells on micropatterns (25) (26). As cardiomyocytes display LR asymmetries during cardiac looping, and heart looping morphogenesis requires actomyosin activity, this presents the exciting hypothesis that vertebrate heart looping depends on tissue- and cell-intrinsic chirality (9, 27, 28).

To identify novel factors and mechanisms that drive cardiac looping, we have performed forward genetic screens in zebrafish (9, 29–31). In such a screen we identified the *oudegracht* (*oug*) mutant in which cardiac jogging was unaffected while cardiac looping was compromised. We found that a novel loss-of-function allele for *tbx5a*, one of the two zebrafish paralogues of *Tbx5*, was responsible for the cardiac looping defect in *oug* mutants. While *tbx5a* has been implicated in cardiac morphogenesis, a link to intrinsic heart looping morphogenesis has not been established yet. Tbx5 is a transcription factor which acts as a master regulator of cardiac development, with established roles in cardiomyocyte differentiation, conduction system development and septation across vertebrates, including humans (32, 33). To gain a better understanding of cardiac looping we performed live two-photon confocal imaging in wild type and *oug* mutant embryos and mapped cardiomyocyte behavior at a single cell level. Our study establishes that during looping, cardiomyocytes in the forming ventricle and atrium actually rearrange in opposite directions towards the outer curvatures of the chambers. Hence, the ventricle and the atrium undergo asymmetric, opposite rotational movements around the atrioventricular canal, effectively transmitting a twisting transformation to the heart tube, a process which we show to be defective in *tbx5a*^*-/-*^ zebrafish mutants. To address which cellular processes exert a regulatory role in this major cellular rearrangement, we manipulated cardiac looping by chemical treatment or *ex vivo* culture and analyzed single cell behavior during heart morphogenesis. Finally, rescue of the *tbx5a*^*-/-*^ cardiac phenotype in a *tbx2b*^*-/-*^ background establishes that the intrinsic looping morphogenesis relies on correct genetic patterning during cardiac development.

## Results

### Tbx5a is required for cardiac looping and patterning

We have performed several forward genetic screens to identify genes that regulate L/R patterning and heart looping morphogenesis (9, 29–31). In short, embryos were screened around 28 hpf for correct formation and asymmetry of the cardiac tube, and at 50 hpf to assess cardiac looping. In one of these screens the recessive and lethal *oudegracht* (*oug*) mutation was identified, named after the stretched S-shaped canal in the city of Utrecht (NL). The *oug* mutants displayed cardiac edema, defective cardiac looping at 50 hpf (Fig.1A-C) and reduced heartbeat rate (not shown). LR patterning was unaffected in *oug* embryos since the direction of cardiac jogging was predominantly leftward and the laterality of the visceral organs was not affected (Fig. 1C). Morphologically, *oug* mutants grow normally, though importantly they lack development of the pectoral fin buds (Fig.1B). Using positional cloning and direct sequencing, we determined that *oug* mutants carry a point mutation resulting in a premature truncation of the Tbx5a transcription factor (Figure 1D-F; ENSDARG00000024894). We carried out a complementation test with a previously identified *tbx5a* mutant allele, *heartstrings (hst)* which was also reported to display cardiac looping and fin bud formation defects (34). Crossing of heterozygous *oug* and *hst* carriers to each other failed to rescue both these phenotypes, and thus confirmed that *oug* is a *tbx5a* allele (Fig.1G).

**Figure 1.**
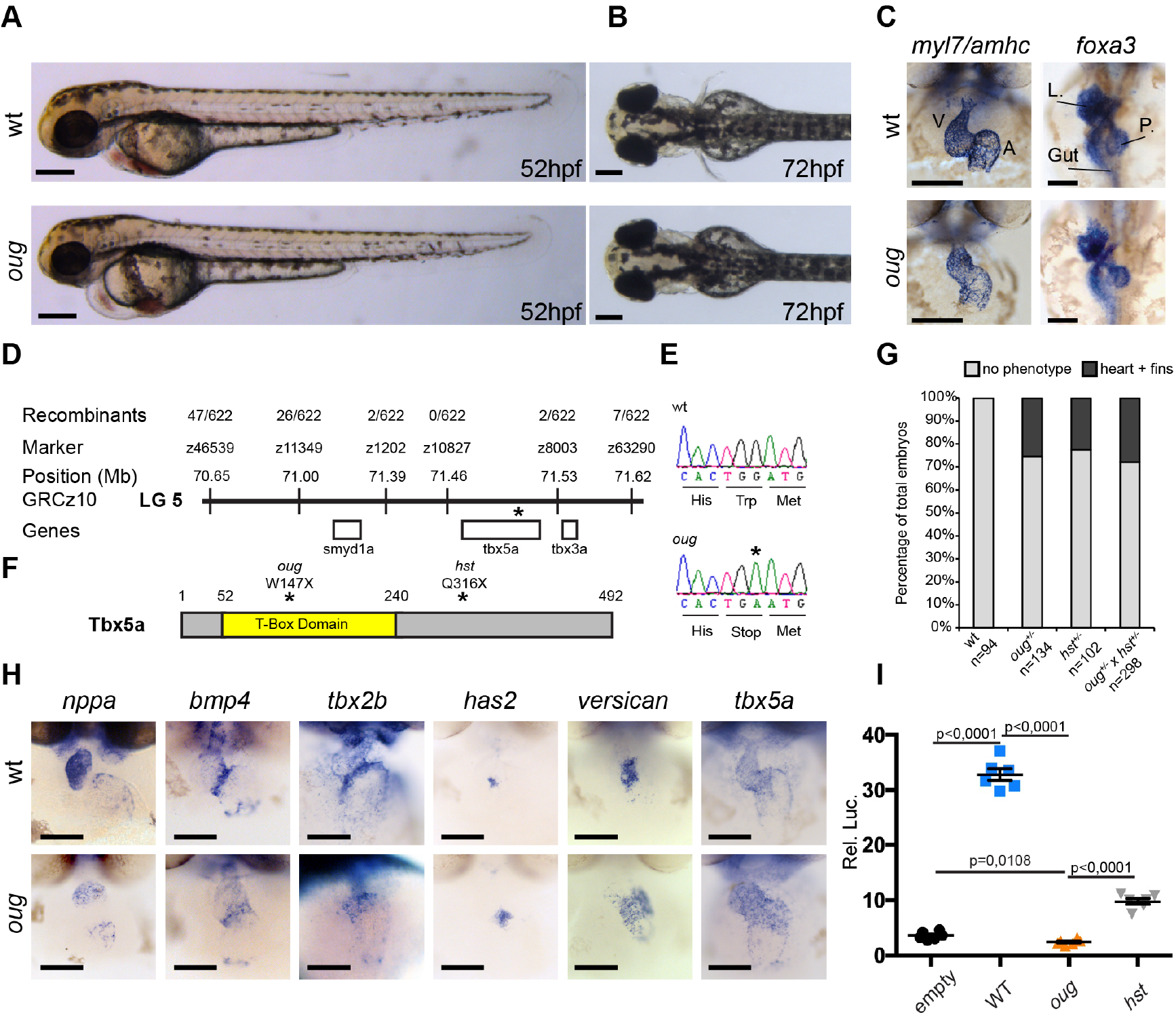
The *oudegracht (oug*) mutant carries a *tbx5a* null allele and displays defective cardiac looping. **(A)** Lateral view of wt and *oug* mutant embryos at 52 hpf. Note the cardiac edema in *oug*. **(B)** At 72 hpf dorsal observation of *oug* mutant embryos reveals absence lateral fins. **(C)** 2 dpf *oug* mutant embryos display defective cardiac looping but normal asymmetric positioning of the internal organs. L, liver; P, pancreas **(D)** Mapping and genomic position of the *oug* mutation (indicated by the asterix). **(E)** A single nucleotide substitution in *tbx5a* (G to A) resulting in a tryptophan (Trp; TGG) to stop (TGA) mutation segregates with the *oug* phenotype. **(F)** Tbx5a is truncated at amino acid 147 in *oug*, in its T-Box domain. The *hst* allele (Q316X;(34)) is included for comparison **(G)** Complementation test. Outcross of *oug*^+/−^ to *hst*^+/−^ fails to complement the *oug* cardiac and pectoral fin bud phenotype. **(H)** Gene patterning is affected in *oug* hearts at 2dpf. Expression of *nppa* is reduced in the cardiac chambers while expression of *bmp4* and *tbx2b* is expanded in the AV canal. Cardiac cushion markers *has2* and *versican* also show expanded expression domains. Transcripts for *tbx5a* are detected in wt and *oug* mutants. **(I)** Luciferase assay establishes that *oug* retains virtually no activity. Shown values are mean values +/− SEM. Statistics: 2-tailed unpaired Student’s t-test. Scale bars: 100 μm.

Surprisingly, when we analyzed the patterning of the heart in chamber (working) and AV canal (non-working) myocardium we noticed a striking difference between the *oug/tbx5a* and the reported *hst/tbx5a* phenotype. In situ hybridization (ISH) using markers for the AV canal and chamber myocardium revealed a strong reduction in chamber specification (*nppa*, Fig.1H) while AV canal specification was expanded as revealed by expanded domains of expression for *bmp4* and *tbx2b* (Fig.1H), which contrasted a previous observation that AV canal specification was lost in *hst/tbx5a* mutant hearts (34). In accordance with our observations on the AV canal myocardium, we also detected increased expression for the AV endocardial markers *has2* and *versican* (Fig.1H).

The *oug/tbx5a* allele (hereafter, and throughout the manuscript referred to as *oug*) truncates Tbx5a at amino acid 147 (out of 492; Fig.1F), resulting in the loss of approximately 50% its DNA-binding T-Box domain, which is crucial for its function (35). This is not the case for the *hst/tbx5a* allele, which does not affect T-Box domain (Fig.1F).

To address whether the striking difference in AV canal specification could be due to differences in the effects of the *oug* and *hst* alleles on Tbx5a activity we carried out an *in vitro* test for Tbx5a activity (Fig.1I). Our results show that while the *oug* allele causes an almost complete loss of Tbx5a activity the *hst* allele retained a significantly higher capacity to induce luciferase expression (Fig.1I). Hence, the defect in cardiac gene patterning and accompanying failure to complete cardiac looping in *oug* mutant embryos are the result of loss of Tbx5a function.

### Time-lapse imaging reveals twisting of the chambers around the AV canal

Cardiac looping in zebrafish can be observed from 30 hpf and is considered to be completed, including chamber ballooning, at around 55 hpf. During this process the heart tube seemingly undergoes flat bending (or planar buckling) along its anterior-posterior axis (Supplementary Figure S1). To get more insight into this transformation we followed the movements of individual cardiomyocytes from 28 hpf to 42 hpf (Fig.2A) in hearts in which cardiac contractions were suppressed (36). At this early stage, the embryonic zebrafish heart displays normal heart morphogenesis in the absence of heartbeat (9). Individual cardiomyocytes were tracked (Fig.2B) and the start and end point of each trace was used to obtain the individual track displacement (described in the Materials and Methods section), hence quantifying the displacement of each tracked cardiomyocyte and representing it as a vector (Fig.2C). We focused on cellular movements pertaining to the cardiac chambers: based on the starting location at the beginning of their corresponding track, cells were categorized in four regions: ventral or dorsal half of the ventricle (VV and VD), and ventral or dorsal half of the atrium (AV and AD; Fig.2D-H). We observed that while cells within a specific region displayed coherent movements, there were large differences between these four regions. Most strikingly, the vectors in the ventral and dorsal sides of the atrium pointed in opposite directions (Fig.2E-F). If planar buckling was a major contributor to the transformation, the expected displacement vectors for the dorsal and ventral sides of each chamber would be similar. Instead, these movements in opposite directions suggested that the atrium rotates during cardiac looping. To address this further, approximately 30 cells per cardiac region were analyzed in three distinct time-lapse movies per condition. To calculate the angle of cellular displacement we defined the longitudinal axis of the heart at the start of the time-lapse (Fig. 2A;I) and we recorded and quantified the angle of individual displacement vectors with respect to this axis (Fig. 2I,J; Supplementary Figure S2; detail of the calculation in Material and Methods section). This method gave consistent results for the different hearts that were analyzed. Most importantly, we found that in the atria of these hearts, the cardiomyocytes in the ventral half displayed positive displacement angles (θ) while cardiomyocytes in the dorsal half displayed negative displacement angles (Fig. 2I,J). This confirms that within the atrium, cells belonging to the ventral and dorsal sides of the heart tube, respectively, concertedly move in opposite directions during cardiac looping, hence supporting a chamber-specific rotational movement. For the ventricle we also observed a negative correlation between the displacement angles of the cardiomyocytes in either the dorsal or ventral sides. The main difference with the atrium, however, was that while the ventricle displayed a clockwise rotation as viewed from the OFT, the atrium displayed an anticlockwise rotation. The strong difference observed in migration directionality between the cardiomyocytes located ventral and dorsal suggests that simple planar buckling only contributes moderately to cardiac looping. Instead, these results suggest that the heart tube undergoes a twisting transformation, with a “hinge” between the 2 oppositely rotating chambers around the AV canal.

**Figure 2.**
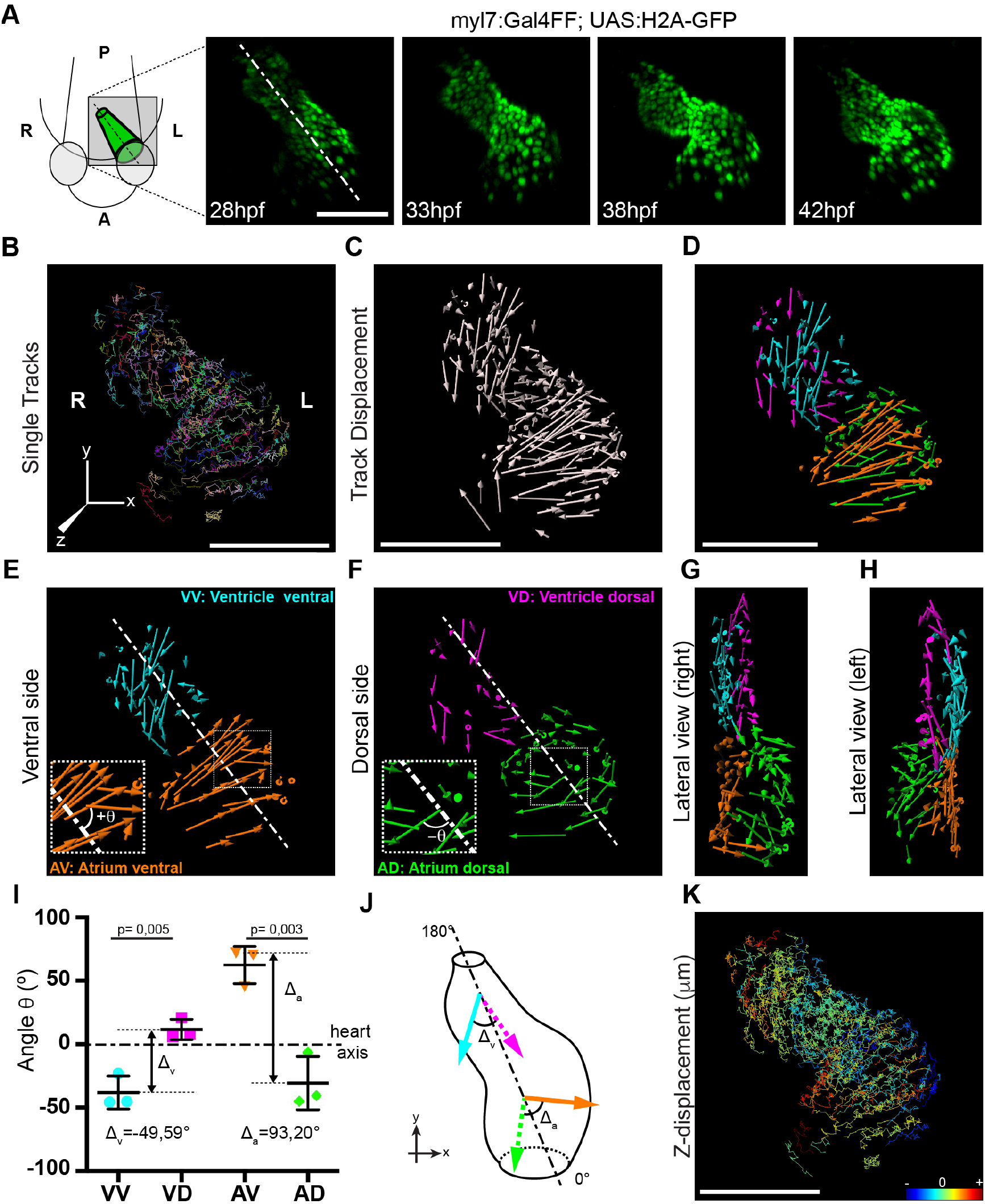
Cardiac looping is accompanied by distinct rotation of the cardiac chambers. **(A)** Time-lapse imaging is carried out on *tg(myl7:Gal4FF; UAS:H2A-GFP)* embryos. Dashed line represents the heart tube axis at 28 hpf. One representative heart is shown. **(B)** Total tracks (Ventral View). Each track is colour-coded and is assigned an ID number. **(C)** Track displacement vectors for each single trace. **(D)** Track displacement vectors to be analyzed are selected, categorized by visual inspection and colour-labeled accordingly **(E)** Detail of the track displacement vectors on the ventral cardiac side. **(F)** Detail of the track displacement vectors on the dorsal cardiac side. Comprehensive visualization of the differences in cellular displacement is achieved. **(G)** and **(H)** Lateral views from right (G) and left (H) of the selected tracks. **(I,J)** Quantification of the mean displacement angles per heart region (I) and representation on schematized zebrafish heart (J). **(K)** Colour-coding of total tracks according to their displacement on the Z axis. Statistics for (I): Each of the 3 data points represents 24<n<40 measurements (see Supplemental Figure S2). Statistical analysis: 2-tailed unpaired Student’s t-test; values reported are mean values +/− SEM. Scale bars: 100 μm.

To gain more insight into this process, we also visualized the cardiomyocyte tracks according to their Z-displacement (Fig. 2K). Regionalized concerted cellular movements were also apparent here, especially in the atrium where distances travelled by the cell during morphogenesis are most pronounced. We could detect that cells at the atrial outer curvature (OC) display are retreating (negative displacement in Z; traces with blue tones), which is compatible with a re-localization from the ventral to the dorsal side of the heart tube. As cells at the atrial inner curvature (IC) behaved in the opposite manner (positive Z-displacement, red tone traces), this corroborates an anticlockwise rotation of the atrium with respect to the cardiac axis (defined in Fig. 2A,I,J), taking the OFT as view point. In the ventricle, although visualization is more difficult due to the higher compactness of the tissue, we observed traces compatible with a clockwise rotation of the chamber. Together, these results indicate that during the early phase of heart looping the cardiac chambers twist around the AV canal boundary in opposing directions.

### Lineage tracing of left and right cardiac fields reveals torsion of the cardiac tube

During linear heart tube formation the cardiac disc rotates in a clockwise direction (from a dorsal view), while at the same time invagination of the right- and posterior sides results in a 3-dimensional cone (7, 37–39). As a consequence of this rotation and folding, the cardiomyocytes originating from the left cardiac field form the dorsal side of the tube, while cells originating from the right cardiac field form the ventral side at approximately 24 hpf (16). A model has been proposed in which this clockwise rotation is followed by a counter clockwise rotation just before or during looping, which would restore the original left-right orientation of the cardiac cells (7). This two-rotation model would not be compatible with our observations from the cell tracking of ventricular cardiomyocytes. In an attempt to resolve this, we generated a new transgenic line that would allow an accurate tracing of cells derived from the left and right cardiac fields. The transgenic line, referred to as *tg(0.2Intr1spaw:eGFP)* (Supplementary Figure S3) was made by using a highly conserved 0.2 kb sequence in the first intron of the Nodal-related gene *spaw*, which acts as an asymmetric enhancer (ASE;(40, 41)). This ASE sequence drives GFP expression in the left lateral plate mesoderm (LPM) during somatogenesis. While *spaw* mRNA is no longer detectable in the left heart field beyond 30 hpf, the stability of the fluorescent protein allows us to follow left-derived GFP positive cells up to 2 dpf (Supplementary Figure S3). This line could therefore be used in combination with a *myl7* fluorescent reporter to trace cells originating from the left and right cardiac fields during cardiac looping stages and address how these cells behave during cardiac looping morphogenesis.

We first wanted to test whether we could confirm the clockwise rotation during linear heart tube formation, which results in left-derived cells occupying the dorsal side and right-originating cells occupying the ventral side of the tube (37, 38). Indeed, this clockwise rotation is also observed *in vivo*, in *tg(myl7:Gal4FF; UAS:RFP; 0.2Intr1spaw:eGFP)* zebrafish embryos (Fig.3A, A’). We then proceeded to use these transgenic lines to analyze the localization of the left- and right-originating cells in the looped heart. Interestingly, at this stage, left-originating cells seem to localize mainly to what now is the ventral side of the heart tube (Fig.3B,B’’). In addition, we observed left-originating cells at the outer curvatures of both chambers (Fig.3B’), reaching, especially visible in the ventricle, the dorsal side of the heart (Fig.3B’’, arrowheads). Concomitantly, the inner curvature of the atrium is only RFP-positive, indicating the right origin of these cardiomyocytes (Fig.3B’). To confirm these observations, we generated an additional reporter line in which the regulatory sequences of the *lefty2* gene drive expression of Gal4FF (42), referred to as *tg(lft2BAC:Gal4FF)* (Supplementary Figure S4A). This line, when combined with a UAS fluorescent reporter line, recapitulated endogenous *lefty2* expression (Supplementary Figure S4B). Analysis of the localization of the left- and right-originating cells in the looped heart in *tg(lft2BAC:Gal4FF)* by fluorescence immunolabeling (Supplementary Figure S5) corroborated our results obtained with *tg(0.2Intr1spaw:eGFP)*. Together, these observations are consistent with those from our time-lapse imaging and cell tracing. Furthermore, they confirm our conclusion that cardiac chambers twist around the AV canal in opposing directions resulting in torsion of the heart tube.

**Figure 3.**
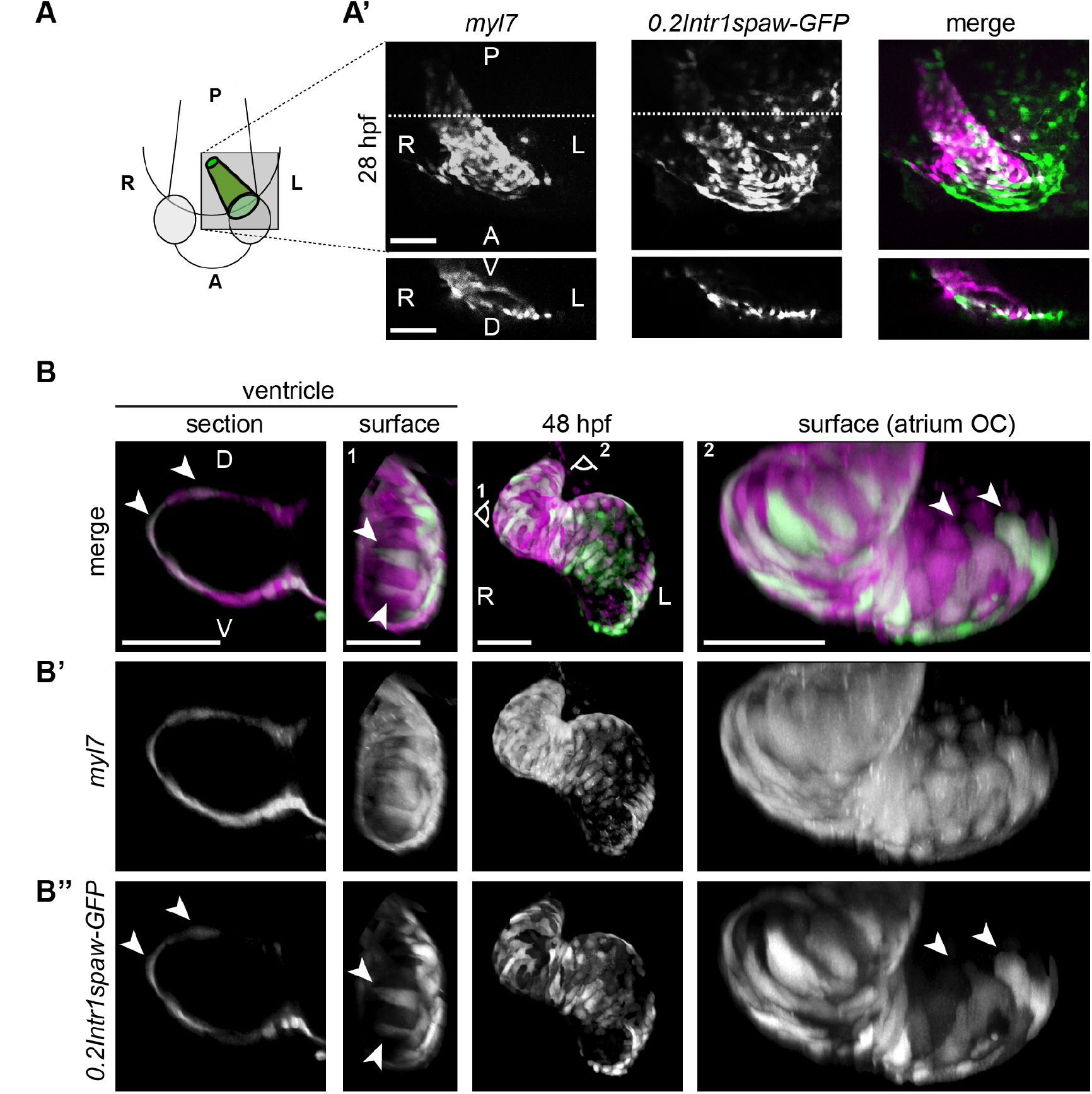
Origin and final positioning of left- and right-originating cardiomyocytes during cardiac looping. **(A)** At 28 hpf, as cardiac jogging towards the anterior left side of the embryo is completed, **(A’)** the *tg(0.2Intr1spaw:eGFP)* labels cardiomyocytes localizing to the dorsal side of the cardiac tube (section). **(B, B’, B’’)** By 48 hpf cardiac looping morphogenesis is accompanied by displacement in opposite directions of left-originating cardiomyocytes (green, B’’) towards the outer curvatures of the ventricle and the atrium (arrowheads in the section and surface view panels). Scale bars: 50 μm.

### Tbx5a is required for the twisting of the cardiac chambers

To address the role of Tbx5a in the observed twisting of the cardiac chambers we time-lapsed and analyzed cardiomyocyte displacements in *oug* mutant embryos in the same manner as we did for siblings using the *tg(myl7:Gal4FF; UAS:H2A-GFP)* line (Fig.4A-I). Strikingly, we observed that the directions of displacement of cardiomyocytes in the ventral and dorsal sides of both chambers are relatively similar (Fig.4B-H). Indeed, when we compared the angles of displacement (θ) of cardiomyocytes in the ventral and dorsal sides of the cardiac chambers we observed no significant differences (Fig.4H,I Supplementary Figure S6). Furthermore, important cellular movements in the Z-axis were not evident from the analysis (Fig. 4J). This suggests that the ventral and dorsal sides of the *oug* mutant heart move relatively concertedly between 28 hpf and 42 hpf, which is compatible with bending of the cardiac tube. Hence, the twisting, as observed in wild type hearts was largely nonexistent in *oug* mutant hearts, limiting the morphogenesis process to moderate planar buckling of the heart tube.

**Figure 4.**
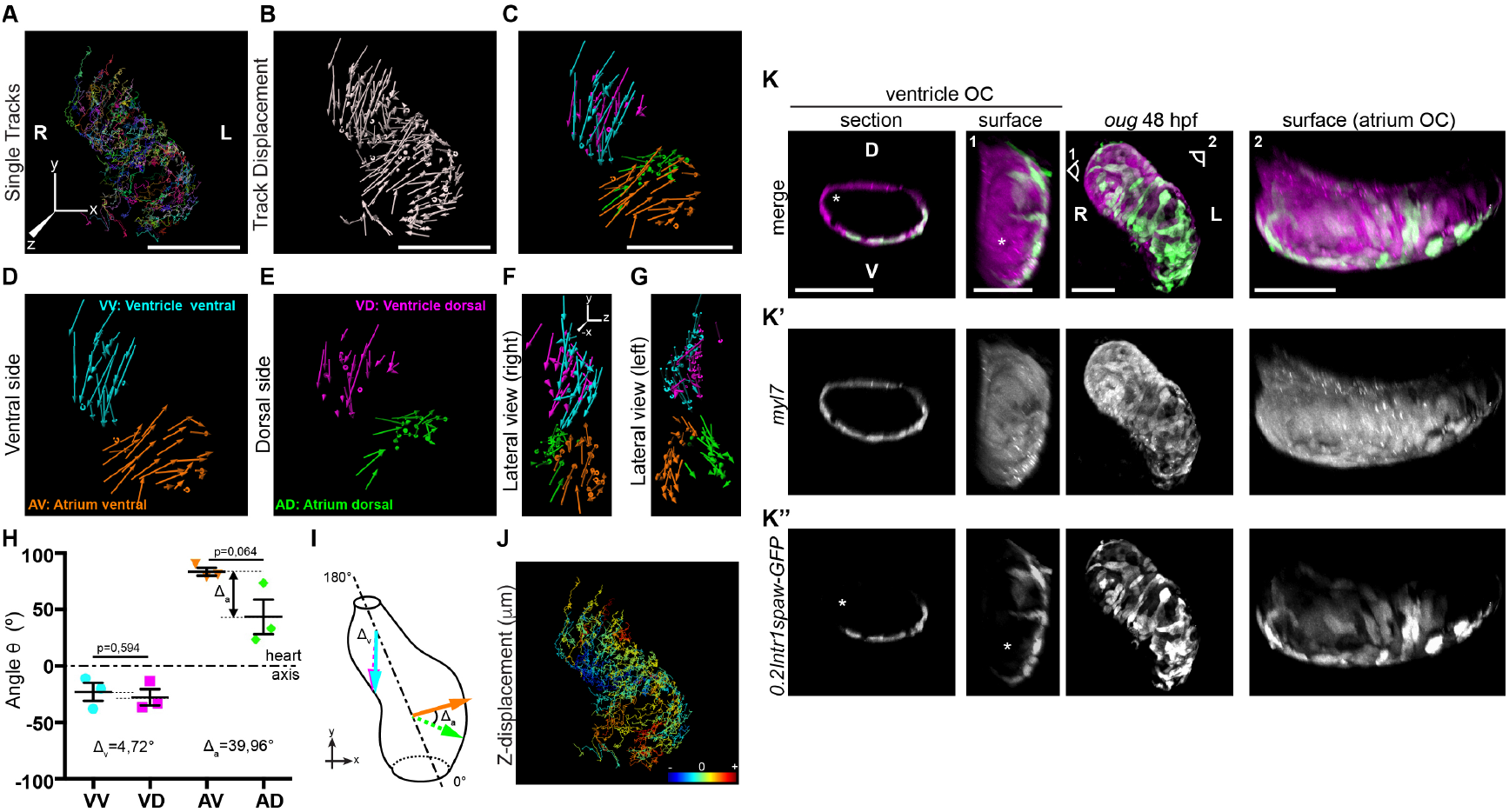
Cardiac looping is defective in *oug* mutants due to absence of asymmetric rotation of the cardiac chambers. **(A)** Total tracks (Ventral View) obtained from a time-lapse movie of cardiac looping in an *oug* mutant. Each track is colour-coded and is assigned an ID number. **(B)** Track displacement vectors for each trace drawn in (A). **(C)** Track displacement vectors to be analyzed are selected, categorized by visual inspection and colour-labeled accordingly. **(D)** Detail of the track displacement vectors on the ventral cardiac side and **(E)** detail of the track displacement vectors on the dorsal cardiac side. Comparison with sibling (Fig.2) reveals differences in concerted cellular movements **(F)** and **(G)** Lateral views from right (F) and left (G) of the selected tracks. **(H; I)** Quantification of the average displacement angles per heart region (H) and representation on schematized zebrafish heart (I). **(J)** Colour-coding of total tracks according to their displacement on the Z axis. **(K, K’, K’’)** At 48 hpf the *oug* mutant heart tube fails to display any constriction at the AV canal and left-originating cardiomyocytes (green; K’’) are not visible at the outer curvatures of the cardiac chambers (asterisk; ventricle). Analysis was carried out in identical manner as for wt hearts presented in Fig. 2. Statistics for (H): Each of the 3 data point represents 16<n<30 measurements (see Supplemental Figure S6) collected on 3 different hearts. 2-tailed unpaired Student’s t-test; horizontal bars: mean value +/− SEM. Scale bars: (A-C): 100 μm; (K): 50 μm.

To address the consequences for the reduced twisting of the chambers on the torsion of the heart tube in *oug* mutants we crossed the *oug* mutation into the *tg(myl7:Gal4FF; UAS:RFP; 0.2Intr1spaw:eGFP)*. Contrary to observations in wild type hearts, we observed that the outer curvature of both the ventricle and atrium in *oug* mutant hearts are largely devoid of left-originating GFP+ cells (Figure 4K). In transversal sections of the ventricle, left-originating cells remain largely localized to the ventral side of the heart tube (Figure 4K-K’’). The domain occupied by left originating-cells remained virtually unchanged when compared to the situation at the end of cardiac jogging, corroborating a lack of twisting and the absence of torsion in hearts lacking Tbx5a.

### A tissue intrinsic mechanism, and not cell addition to the embryonic cardiac poles, is required for torsion of the heart tube

Next, we wanted to address which mechanisms could be driving the observed opposite twisting of the chamber around the AV canal during heart looping. During mouse heart morphogenesis, asymmetric contributions at the poles drive a helical rotation of the tube (43). Although the zebrafish heart does not form a helix, we considered that the opposite chamber rotation could be driven by a similar mechanism. Previous work has demonstrated that also in zebrafish cells from the second heart field (SHF) are added to the poles of the heart tube concomitantly with cardiac looping (44–46). To test whether cardiomyocyte addition from the SHF is required for the correct progression of cardiac looping, we abolished it in two independent manners prior to the onset of cardiac looping: 1) by treating embryos with the FGF inhibitor SU5402 (44) and 2) by explanting linear heart tubes and culturing them *ex vivo* for 24h, as previously described (9). In SU5402–treated whole embryos, we observed robust cardiac looping in the majority of the hearts, although the cardiac tube is clearly reduced in size and general development is affected (Figure 5A). In explanted cultured *tg(lft2BAC:Gal4FF; UAS:RFP; myl7:GFP)* hearts (Figure 5B,C), we also observed convincing cardiac looping (Figure 5C, upper panels). The use of the *lft2* reporter allowed us to observe that left-originating cardiomyocytes locate to the outer curvatures of the ventricle and atrium, thus indicating that heart tubes *ex vivo* not only retained their capacity to loop dextrally (9), but also that the cardiac torsion was still occurring. We also exposed explanted heart tubes to SU5402 during culture and still observed satisfactory looping morphogenesis and localization of left-originating cells to the outer curvature of the ventricle and atrium (Figure 5C, lower panels).

**Figure 5.**
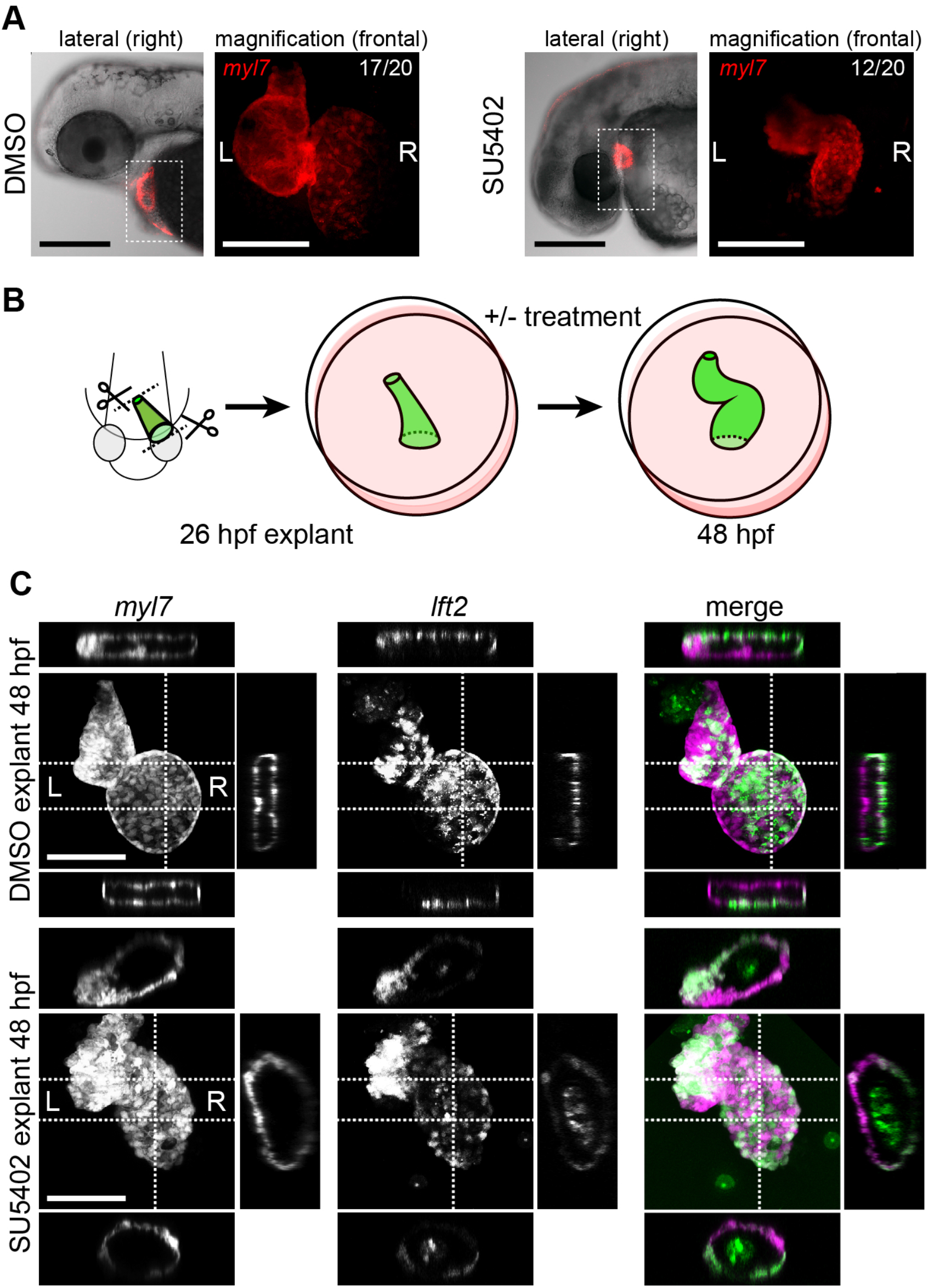
Chemical and physical suppression of cell addition to the heart tube do not affect proper completion of cardiac looping. Representative DMSO Control and SU5402-treated embryos and explanted hearts are shown. **(A)** Treatment between 1dpf and 2dpf with SU5402 affects overall size of the looping heart tube but its capacity to complete cardiac looping. **(B)** Heart explant procedure: as cardiac jogging is completed (26 hpf) heart tubes are explanted and put into culture for approximately 24 hpf during which chemical treatments can be carried out. At 48 hpf the hearts are imaged. **(C)** Heart tubes explanted at 26 hpf and subsequently cultured in liquid medium for 24h display normal formation of a ventricle, atrium and atrioventricular canal. The *lft2* reporter allows visualization of left-originating cells at the outer curvature of both ventricle and atrium, in control (DMSO) and treatment (SU5402) conditions. Scale bars: 100 μm.

Consistent with our observation that addition of SHF cells to the poles of the heart tube is dispensable for opposite chamber rotation and cardiac looping, we observed no changes in cardiomyocyte numbers in the ventricle of *oug* mutants (Figure 6A,B). To reject the possibility that the looping phenotype displayed by *oug* mutants is not secondary to fluid pressure caused by the cardiac edema appearing by 2 dpf, we explanted *oug tg(myl7:Gal4FF; UAS:RFP; 0.2Intr1spaw:eGFP)* heart tubes at 28 hpf. Indeed, after 24h *in vitro* culturing, *oug* mutant hearts failed to loop, indicating that the morphogenesis defect was not related to extrinsic cues in the embryo (Figure 6C). From the above results we conclude that cardiomyocyte addition from the SHF is dispensable for cardiac torsion and helical looping.

**Figure 6.**
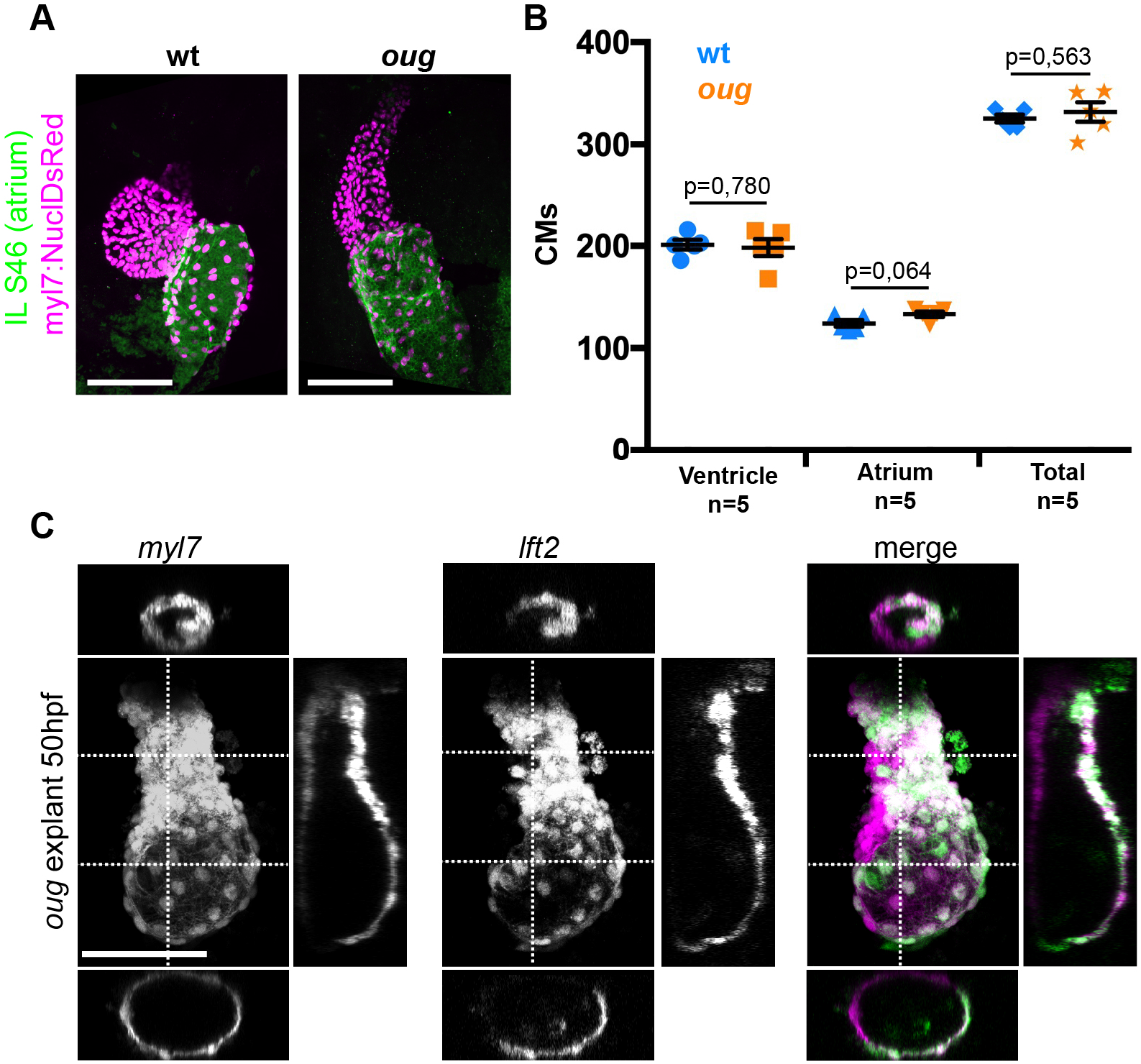
Defective looping in *oug mutants* is not due to cardiomyocyte number or embryonic cues extrinsic to the heart. **(A)** Immunofluorescence with atrium-specific S46 antibody allows distinction of the cardiac chambers. **(B)** Quantification of ventricular and atrial cardiomyocytes in wt and *oug* mutant embryos at 2dpf. **(C)** Explanting *oug* mutant hearts and culturing them *in vitro, ex-embryo* does not rescue defective looping. Statistical analysis: 2-tailed, non-paired Student’s t-test. Horizontal bars: mean value +/−SEM. Scale bars: 100 μm.

### Reduced anisotropic growth in *oug* cardiomyocytes

Epithelial remodeling is an important driver for asymmetric rotation of the Drosophila gut tube or looping of the chick midgut and heart tube (19, 28, 47). In the zebrafish heart tube epithelial remodeling, as observed by changes in cardiomyocyte shape and cell boundaries, occurs during looping morphogenesis as well (27, 48, 49). Hence, we next proceeded by assessing the shape of ventricular cardiomyocytes between 30 hpf and 42 hpf (Fig. 7A). We focused on the outer curvature of the ventricle, as this is a region that undergoes clearly observable transformation (This study, (27)). Indeed, we could determine that the progression of the left-originating cardiomyocytes is concomitant to anisotropic growth of these cardiomyocytes, which results in a reduced roundness (Fig. 7B). Analysis of the positioning of left-(green) and right-(red) originating cardiomyocytes at the OC of the ventricle confirmed this change in cell shape (Fig.7C-D), possibly suggesting involvement of cell intercalation. In *oug* mutant embryos, we observed that ventricular cells retain their higher cell roundness throughout the analysis window and display a much straighter left/right boundary at the ventricular outer curvature. We therefore conclude that our results are consistent with the proposed model in which tissue intrinsic properties drive opposite chamber rotation and cardiac looping (9, 27) and indicate that Tbx5a activity is required for this to occur.

**Figure 7.**
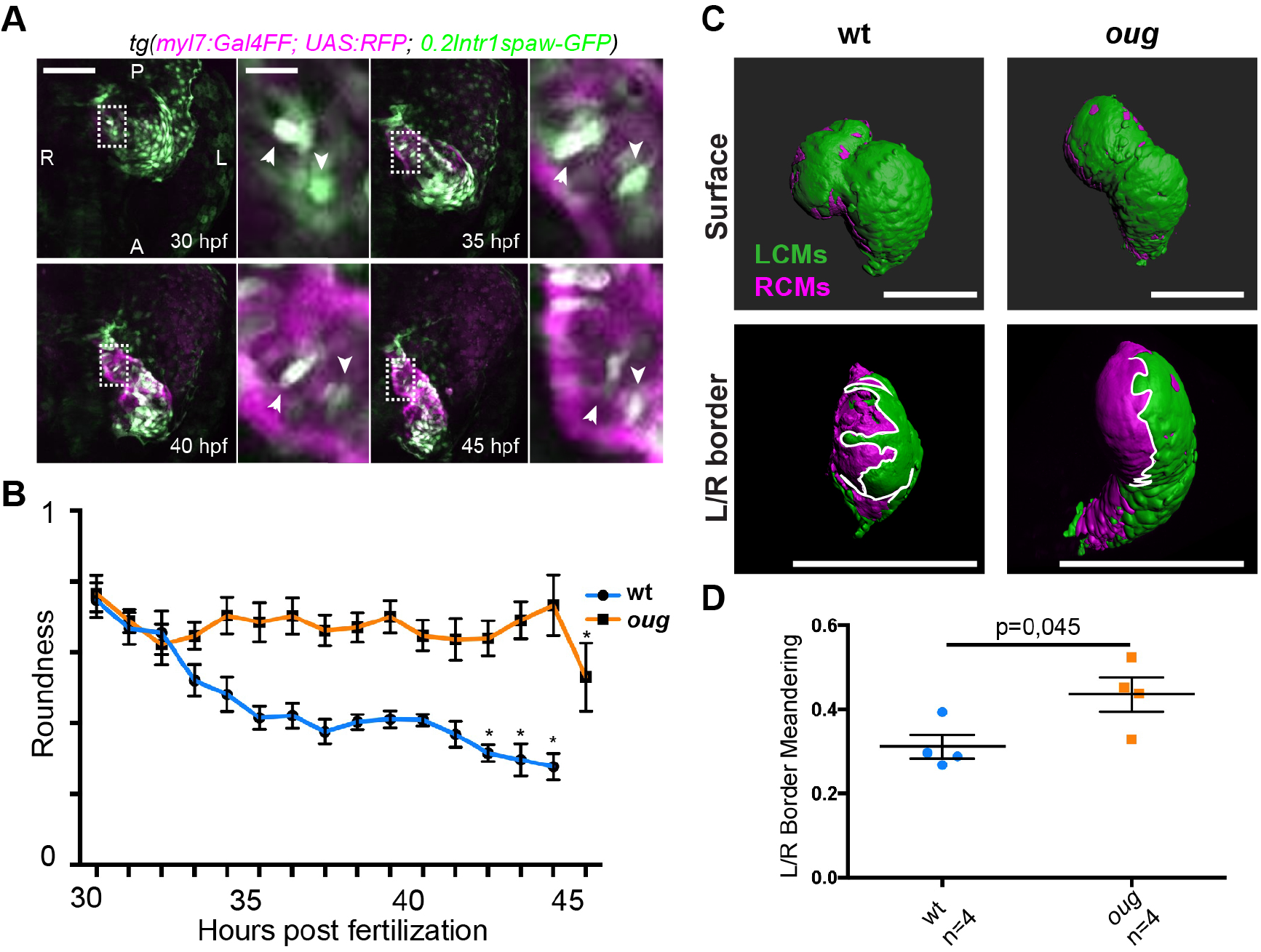
Anisotropic cell shape changes at the outer curvature accompany cardiac looping. **(A)** Time-lapse imaging reveals that cardiac looping is accompanied by anisotropic cellular shape changes in the outer curvature of the ventricle. **(B)** Quantification of cell roundness as observed in (A) and comparison between values for wt and *oug* mutants. **(C)** Surface rendering of *tg(myl7:Gal4FF; UAS:RFP; 0.2Intr1spaw-GFP)* in 48 hpf hearts allows clear definition of a boundary between Left-originating cardiomyocytes (LCMs, green) and right-originating cardiomyocytes (RCMs, red), which allows calculation of the meandering index of the left/right boundary at the outer curvature of the ventricle. **(D)** Quantification of the meandering index is indicative of the level of anisotropic growth in wt and *oug* mutant hearts. Horizontal bars: mean value +/− SEM. Scale bars: (A) 100 μm, magnification 20 μm; (C) 100 μm. Data points in (B): for all points 5<n<9 unless *: n=2. Statistical analysis in (D): 2-tailed, non-paired Student’s t-test.

### Cardiac looping is reestablished in Tbx5a-defective hearts by suppression of Tbx2b activity

AV canal versus chamber specification is tightly regulated by a balance in gene activation and repression by Tbx5 and Tbx2, respectively (50, 51), reviewed in (52). As we observed an expansion of *tbx2b* expression in *oug* mutant hearts (Fig.1D), we first tested whether the myocardial patterning defect in *oug* mutants could be rescued by reducing Tbx2b activity. To do so, we used the *tbx2b* mutant *from beyond* (*fby*) (53). Analysis of cardiac markers by ISH and transgenic reporters revealed that *fby/tbx2b*^*-/-*^ embryos display robust cardiac looping and a properly patterned heart (Supplementary Figure S7 and Fig.8). In *tbx5a*^*-/-*^;*tbx2b*^*-/-*^ (*oug/fby*) double mutant background, ISH indicated rescue of the constriction at the AV canal (Fig.8A), re-establishment of *nppa* expression in the cardiac chambers, while *bmp4* expression remained similar to that of *tbx5a-/-* hearts (Supplementary Figure S7). Analysis of *tg(nppaBAC:mCitrine) in vivo* confirmed the rescue of *nppa* expression in the atrium OC of *tbx5a*^*-/-*^;*tbx2b*^*-/-*^ double mutants, which was absent in *oug* embryos (Fig.8B). Next, we investigated how the rescue in cardiac patterning affects heart looping morphogenesis. Along with the reestablishment of myocardial patterning we also observed a significant rescue of the looping phenotype by measuring the looping angle (Fig.8C). Consistently with these observations, analysis of *tg(myl7:Gal4FF; UAS:RFP; 0.2Intr1spaw-GFP)* in *tbx5a*^*-/-*^;*tbx2b*^*-/-*^ embryonic hearts revealed presence of GFP+ left-originating cardiomyocytes at the ventricle OC (Fig.8D’’’), indicating substantial rescue of the twisting of the heart tube. Additionally, we observed that while pectoral fin development was not rescued in *tbx5a*^*-/-*^;*tbx2b*^*-/-*^ double mutants, these fish hardly developed a cardiac edema, as compared to *oug* mutants (Supplementary Figure S8). Altogether, these results indicate that heart looping morphogenesis is the result of proper tissue patterning, and requires a fine balance Tbx5a and Tbx2b activity.

**Figure 8.**
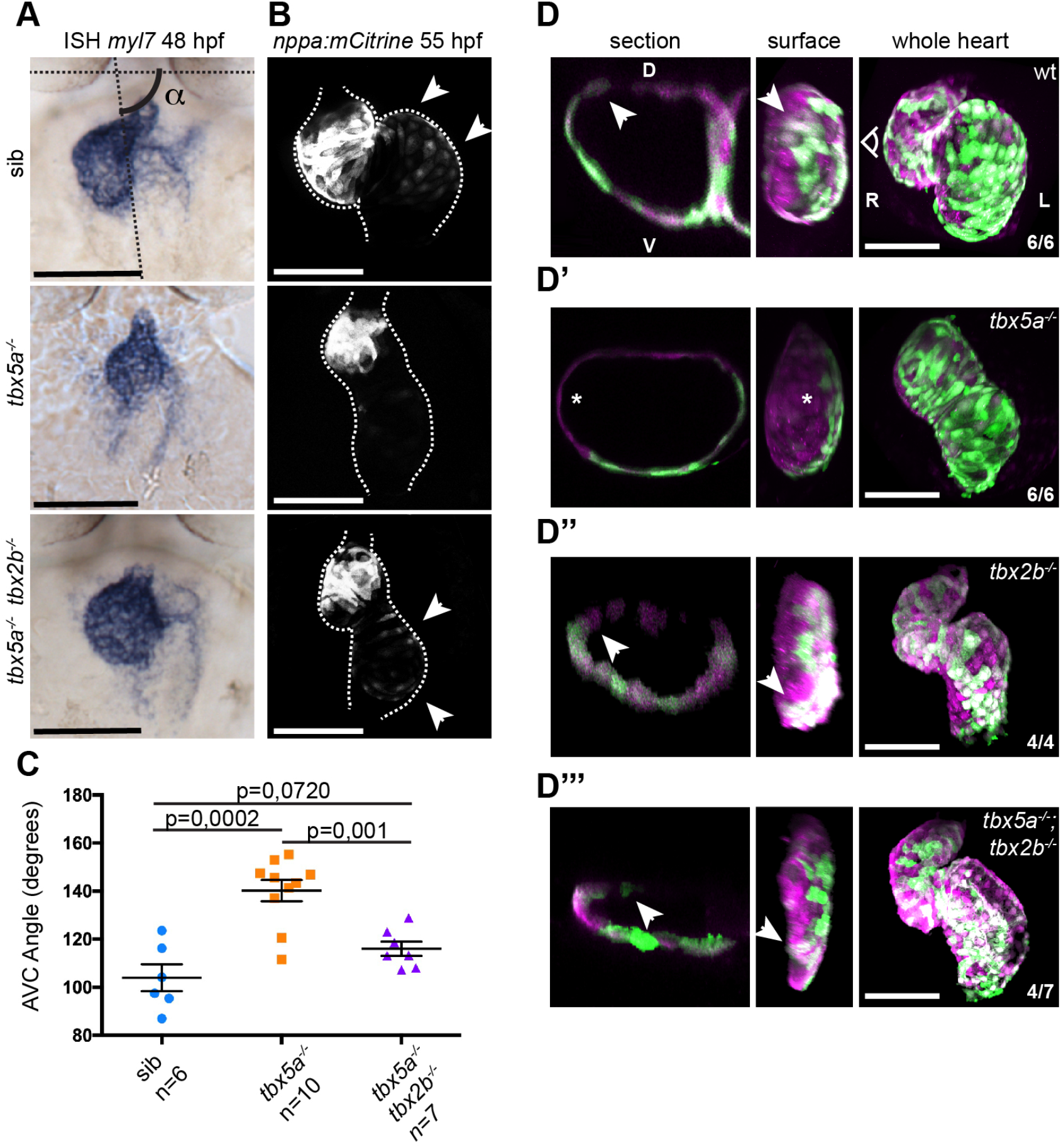
Defective cardiac looping in *oug* mutants is alleviated by simultaneous loss of *tbx2b.* **(A)** ISH for *myl7* at 50 hpf in wild type siblings, *oug* mutants and *tbx5a;tbx2b* double mutants. AV canal angle measurement is exemplified in the sib panel. **(B)** Confocal maximum projections of 2dpf *tg(nppa:mCitrine)* hearts. In the *tbx5a;tbx2b* double mutants, atrial expression of *nppa*, which was lost in *oug* mutants, is re-instated. **(C)** Quantification and comparison of AV canal angles in wild type siblings, *tbx5a* mutants and *tbx5a;tbx2b* double mutants. **(D-D’’’)** 48 hpf *tg(myl7:Gal4FF; UAS:RFP; 0.2Intr1spaw-GFP)* hearts. Wt (D) and *tbx5^-/-^* (D’) are shown for comparison. *tbx2b^-/-^* hearts (D’’) display robust dextral looping and left-originating cardiomyocytes (green) at the ventricle outer curvature, similar to wt (arrowheads in D; Figure 3B-B’’). In double homozygous mutants *tbx5a^-/-^; tbx2b^-/-^* (D’’’) rescue of cardiac looping is observed, accompanied by presence of left-originating cardiomyocytes at the ventricle OC (Compare with D, D’’). Statistical analysis: 2-tailed non-paired Student’s t-test. Horizontal bars: mean value +/− SEM.

## Discussion

Our analysis of cardiac looping is focused on the early phase of this process between 28 hpf and 42 hpf stage, during which time window a distinct S-shaped heart is formed. Very little was known about the cellular behavior during these initial stages of heart looping. Our time-lapse and cell tracing analysis show that cardiomyocytes in each of the 2 chambers move in opposite directions, which results in twisting of the chambers around the AV canal and torsion in the heart tube (Fig. 9). The twisting of the chambers around the AV canal requires proper patterning of the myocardium into chamber and AV canal myocardium, which is regulated by T-box containing transcription factors.

**Figure 9.**
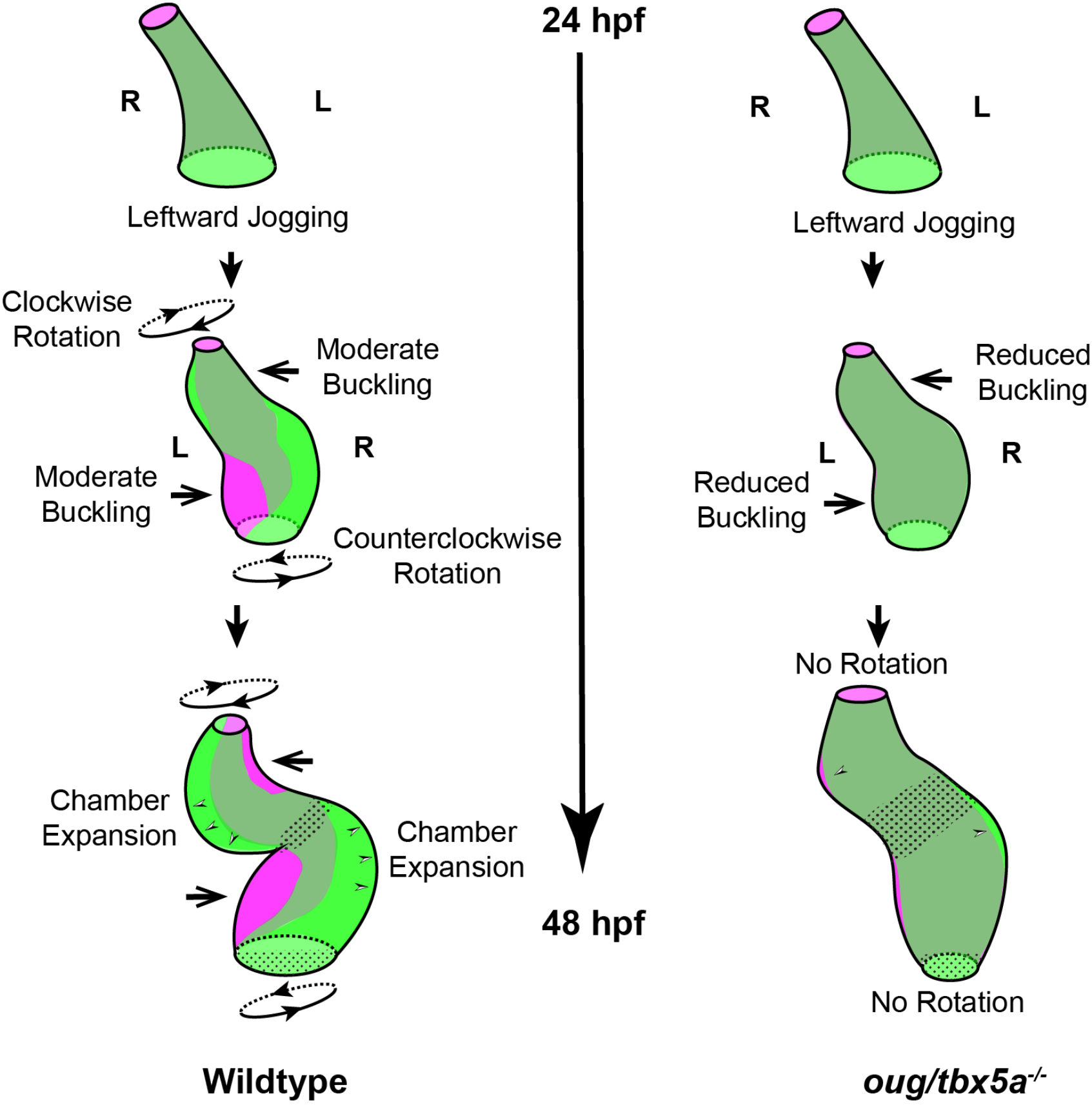
Model for cardiac looping morphogenesis. Viewpoint for describing direction of rotation is always the outflow tract (OFT). In wild type hearts, at the end of cardiac jogging, twisting of the heart tube results in expansion of the space occupied by left-originating cardiomyocytes towards the outer curvatures of both the ventricle and atrium. The resulting torsion of the heart tube is driven by the clockwise rotation of the ventricle and counterclockwise rotation of the atrium, around a fixed hinge, the AV canal. Arrows represent buckling. Bicolor arrowheads at 48 hpf represent chamber expansion. In *oug* hearts, cardiac jogging is completed properly, but progression of cardiac looping is defective. No twisting of the heart tube takes place and reduced buckling and chamber expansion are observed. Defective looping is accompanied by an expansion of the expression domain of *tbx2b* (spotted pattern), especially noticeable at the AV canal (see also Fig.1 and Fig. 7).

In this study we identified a novel *tbx5a* allele, *oug*, which we demonstrated to be a *tbx5a* null allele. Indeed, in *oug* approximately 75% of the gene product is lost, including a large portion of the DNA-binding T-Box domain. While *oug* mutants display an expansion of genes that mark the AV canal, in *hst* mutants the expression of these genes is lost. Based on the *hst* results a model was proposed in which Tbx5a stimulates the expression of *tbx2* in the AV canal (34, 54), which needs to be reconsidered based on the *oug* results. These different outcomes in patterning of the AV canal and chamber myocardium might be explained by the presence of the T-box domain in the previously published *hst* (34), which is only missing regions proposed to affect its subcellular localization (54).

There is a striking resemblance between the rotation in the ventricle during looping as described here and the clockwise rotation that occurs earlier when the cardiac disc transforms into a linear heart tube, which has been described in several studies (7, 38, 39). As a consequence of this first rotation event, the original left-right orientation of the cardiac cells is transformed to a dorsal-ventral orientation. In a previously published study, the authors suggested that after the linear heart tube is formed this dorsal-ventral orientation is transformed back to the original left-right orientation due to a second counter clockwise rotation around its longitudinal axis (7). Although we detected atrial cardiomyocyte movement compatible with this observation (Fig. 2), we did not observe this second rotation when tracing the ventricular cardiomyocytes originating from the left and right lateral plate mesoderm. This difference between the observations might be partially explained by how the left and right cardiac cells were labeled in the two studies. In our study we used stable transgenic lines in which *lefty2* or *spaw* regulatory elements drive left-sided expression of GFP. In the original study by Baker *et al.* (7), a *myl7:Dendra* plasmid was injected at the one- or two-cell stage and embryos were screened before 18 hpf for either left- or right sided expression and analysed at 48 hpf. As we know now, at 18 hpf the *myl7* promoter is only activated in the first heart field (FHF). Cardiomyocytes from the second heart field (SHF) initiate *myl7* expression at a later stage, up to 38 hpf, when these are added to the cardiac poles (44, 45). As a consequence, embryos scored with unilateral *myl7:Dendra* expression at 18 hpf may display expression of Dendra in cardiomyocytes from the originally (18 hpf) non-expressing side when scored at 48 hpf. The gradual activation of *myl7* due to the continuous process of cardiomyocyte differentiation during heart tube morphogenesis limits its use a cell tracing technique.

The clockwise rotation we observed in the ventricle is in the same direction as the rotation that was observed during linear heart tube formation (38). Recently, a clockwise rotation was also described in the OFT of the zebrafish heart at later cardiac looping stages (40-54 hpf) (49). Together, these observations suggest that a clockwise rotation of the cardiac tissue is initiated during linear heart tube formation (20-26 hpf) and that this clockwise rotation continues in the ventricle (28-42 hfp) during looping initiation and continues in the OFT (40-54 hpf) during the late looping stage. In the atrium, however, we describe here a counterclockwise rotation during the early looping phase (28-42 hpf), resulting in a torsion of the heart tube.

During cardiac looping, there is extensive growth of the myocardium. Due to the addition of cells at the poles from the SHF, the number of cardiomyocytes is doubled between 24 and 48 hpf (44). Reduced cell addition from the SHF by inhibiting FGF signaling still allowed looping and twisting of the zebrafish heart tube (Fig. 5). This is different in the mouse heart, where reduced growth due to compromised addition of cells from the SHF results in looping defects (55–57). This may be due to more extensive growth of the murine heart, which extends its length over 4-fold during looping, resulting in a distinct helical shape (43).

Our data builds upon previous work exploring the intrinsic capacity of the heart to loop (9, 28, 58). Corroborating such a model, we observed that the twisting and looping of the heart tube still occurs in conditions in which cell addition is suppressed. We therefore conclude that the early phase of heart looping in zebrafish occurs independently of extrinsic cues (Fig.3). Other examples of tubes that undergo looping morphogenesis due to intrinsic LR asymmetry are the Drosophila genitalia and hindgut (18, 19). For these tubes it is proposed that intrinsic chirality of the epithelial cells drive looping morphogenesis. Like the Drosophila gut tube, the heart tube consists of an epithelial tissue with distinct apical-basal polarity (for references see (16)). During heart looping and chamber ballooning, the epithelium undergoes remodeling, which coincides with regional cell shape changes (27, 48, 49). Interestingly, defective chamber expansion is accompanied in *oug* embryos by failure of the cells at the outer curvature of the ventricle to remodel anisotropically, a process that is regulated by non-canonical Wnt- and PCP-signaling (27). Although regulation by Tbx5 of canonical Wnt ligands is established in limb (59, 60) and lung (61) development, a potential role in controlling cardiac non-canonical Wnt signaling still needs to be explored.

In *oug* mutants *nppa* expression was reduced while *tbx2b* expression was expanded in the AV canal. This was restored in In *tbx5a^-/-^;tbx2b^-/-^* (*oug/fby*) double mutants, which is consistent with the proposed roles of Tbx5 and Tbx2 in patterning the heart in chamber myocardium and primary (e.g. AV canal) myocardium (62). In this respect, it is surprising that no cardiac phenotype was observed in *fby/tbx2b* mutants (Fig.8; Supplementary Figure S7). This could be ascribed to the presence in zebrafish of a second *tbx2* paralogue, *tbx2a*, which is also expressed in the embryonic heart (63). The observed looping defects in *oug* in combination with the observed rescue of cardiac looping in *oug/fby* double mutant supports a model in which cardiac patterning in chamber and AV canal myocardium is an important driver for the intrinsic heart looping morphogenesis.

## Materials and Methods

### Zebrafish Lines

All animal experiments were conducted under the guidelines of the animal welfare committee of the Royal Netherlands Academy of Arts and Sciences (KNAW). Adult zebrafish (Danio rerio) were maintained and embryos raised and staged as previously described (64, 65).

The zebrafish lines used in this study are Tübingen longfin (wild type), *hst/tbx5a* (34), *fby/tbx2b* (53), *tg(myl7:Gal4FF)* (66); *tg(UAS:RFP)* (42); *tg(UAS:H2A-GFP)* (66); *tg(myl7:DsRed)* (67); *tg(mCitrine:nppa)* (68).

### Positional Cloning of *oudegracht/tbx5a*

The *oudegracht/tbx5a*^*hu*6499^ allele was identified in a ENU mutagenesis screen performed as described in (31). The *oudegracht/tbx5a*^*hu6499*^ was mapped using standard simple sequence length polymorphisms (SSLPs)-based meiotic mapping with SSLP primer sequences as pictured in Fig.4. The *oudegracht/tbx5a*^*hu6499*^ mutation introduces a G to A substitution in Exon 4 of *tbx5a* (ENSDARG00000024894) resulting in the introduction of a premature stop codon. The mutation is identified by PCR amplification from genomic DNA using primers FKK106: 5’-GCGCATCAGGTCTGTGAC-3’ and FKK108: 5’-CCAAATACAAGTCCTCAAAGTG-3’ followed by BtscI restriction of the PCR product. The oudegracht/tbx5a^*hu6499*^ mutation removes a BtscI restriction site.

### Generation of transgenic lines

#### tg(lft2BAC:Gal4FF)

The *tg(lft2BAC:Gal4FF)* line was generated by recombineering of bacterial artificial chromosome (BAC) CH211-236P5 as described in (69, 70). A Gal4FF_kan cassette was inserted at the ATG start codon of the first exon of the *lft2* gene (ENSDARG00000044059). Amplification from a pCS2+Gal4FF_kanR plasmid was achieved with primers :
F_LFT2_GAL4FF 5’-cctcagagcttcagtcagtcattcattctttcactggcatcgttagatcaACCATGAAGCTACTGTCTTCTATCGAAC-3’ R_LFT2_NEO 5’-tgtgtgagtgagatcgctgtggtcaaaatgaacagctggatgaacagagcTCAGAAGAACTCGTCAAGA AGGCG-3’

Sequences homologous to the genomic locus in lower case. Recombineering was essentially carried out following the manufacturer’s protocol (Red/ET recombination; Gene Bridges GmbH, Heidelberg, Germany). BAC DNA isolation was carried out using a Midiprep kit (Life Technologies BV, Bleiswijk, The Netherlands). BAC DNA was injected at a concentration of 300 ng/μl in the presence of 0.75U PI-SceI meganuclease (New England Biolabs, Ipswich, MA, USA) in 1-cell stage *tg(UAS:GFP)* or *tg(UAS:RFP)* embryos (both UAS lines:(42)). At 1 dpf, healthy embryos displaying robust *lft2*-specific fluorescence were selected and grown to adulthood. Founder fish (F0) were identified by outcrossing and the progeny (F1) was grown to establish the transgenic line.

#### tg(0.2Intr1spaw:eGFP)

A 228bp conserved sequence located in intron 1 of *spaw* (ENSDARG00000014309) was amplified by PCR using primers FT294 5’-AGTCAAGCATCTCGGGAAGA-3’ and FT295 5’-AGGTCCTGTCAGAGCAGATG-3’. The resulting PCR product was subsequently cloned in the E1b-GFP-Tol2-Gateway construct (Addgene #37846; (71) by Gateway cloning. The resulting construct was co-injected with 25ng/μl Tol2 RNA in 1-cell zebrafish TL embryos. Founder fish (F0) were identified by outcrossing and the progeny (F1) was grown to establish the transgenic line.

### Microinjection of antisense morpholino

The tnnt2a morpholino oligonucleotide targeting the translation start site (5' - CATGTTTGCTCTGATCTGACACGCA - 3') was used to block heart beat (36). We injected approximately 2ng of the oligo morpholino in one-cell stage embryos.

### Chemical treatments

#### SU5402 treatment

Embryos were dechorionated and treated with SU5402 at a concentration of 10 μM in E3 embryo medium from 24 hpf until 48 hpf at 28,5 °C. Control embryos were treated with the corresponding DMSO concentration.

#### Phenylthiourea

Addition of phenylthiourea (PTU) at a concentration of 0.003% (v/v) to the E3 embryonic medium after shield stage (8 hpf) blocked pigmentation for improved confocal analysis.

#### Heart explants

Zebrafish heart tubes were manually dissected from 26 hpf embryos using forceps and placed into supplemented L15 culture medium(15% fetal bovine serum, 0.8mM CaCl2, 50μg/ml penicillin, 0.05 mg/ml streptomycin, 0.05 mg/ml gentomycin) essentially as described in (9). Explants were incubated at 28.5 °C for 24 h and fixed in 4% PFA overnight. Chemical treatment of the explants was carried out in an identical way as for the embryos. Explanted hearts were mounted in Vectashield (Vector Laboratories) before imaging.

#### Immunofluorescent labeling

Zebrafish embryos at the appropriate developmental stage were fixed overnight in 2% paraformaldehyde (PFA) in PBS at 4°C. After washing with 1× PBS–Triton X-100 (0.1%; PBS-T) and blocking in 10% goat serum in 1×PBST (blocking buffer;BB), embryos were incubated overnight at 4°C with rabbit anti-DsRed (1:500 in BB; Clontech 101004), mouse anti-Myh6 antibody (1:200 in BB, Hybridoma bank, S46), or chicken anti-GFP (1:500 in BB, Aves, GFP-1010). After washing in PBST, the embryos were incubated overnight at 4 °C in Cy3-conjugated goat anti-rabbit antibody (1:500 in BB; Jackson Immunoresearch, 111-165-144), Alexa488-conjugated goat anti-mouse (1:500 in BB, Invitrogen, A21133) or Alexa488-conjugated goat-anti-chicken (1:500 in BB; Invitrogen, A11039). Embryos were washed in PBST before imaging.

#### Whole mount mRNA In Situ Hybridization (ISH)

Fixation of the embryos was carried overnight in 4% paraformaldehyde (PFA). Embryos were subsequently stored in methanol (MeOH) at −20°C. Rehydration was carried out in PBST (PBS plus 0.1% Tween-20) and, depending on the stage, embryos were treated with 1 μg ml-1 Proteinase K (Promega) between 1 and 20 min. Embryos were then rinsed in PBST, post-fixed in 4% PFA for 20 min, washed repeatedly in PBST and pre-hybridized for at least 1 h in Hyb-buffer. Digoxigenin-labeled RNA probes were diluted in Hyb-buffer supplemented with transfer RNA (Sigma-Aldrich) and heparin (Sigma-Aldrich), and incubated with the embryos overnight at 70°C. After removal of the probe, embryos were washed stepwise from Hyb-to 2xSSCT, and subsequently from 0.2xSSCT to PBST. Embryos were blocked for at least 1 h at room temperature (RT) in PBST supplemented with sheep serum and BSA before being incubated overnight at 4°C with an anti-digoxygenin antibody (Roche). After removal of the antibody, embryos were washed in PBST before being transferred to TBST. The embryos were subsequently incubated in the dark on a slow rocker in dilutions of Nitro-blue tetrazolium/5-bromo-4-chloro-3-inodyl phosphate (NBT/BCIP; Roche) in TBST. After development of the staining, embryos were washed extensively in PBST and fixed overnight in 4% PFA at 4°C. Before imaging, embryos were cleared in MeOH and mounted in benzylbenzoate:benzylalcohol (2:1). Accession numbers of the genes assayed by ISH: *myl7* (NM_131329), *amhc* (NM_198823), *foxa3* (NM_131299), *nppa* (NM_198800), *tbx2b* (NM_131051), *bmp4* (NM_131342), *has2* (NM_153650), *versican* (NM_001326557) and *tbx5a* (NM_130915).

### *In Vitro tbx5a* activity assay

COS7 cells, grown in 12-well plates in DMEM supplemented with 10% FCS (Gibco-BRL) and glutamine, were transfected using polyethylenimine 25 kDa (PEI, Brunschwick) at a 1:3 ratio (DNA:PEI). Standard transfections were performed using 1.4 μg pGL3basic reporter vector (Promega) containing −638/+70bp r*Nppa* promoter (reporter construct), which was co-transfected with 3 ng phRG-TK Renilla vector (Promega) as normalization control. Zebrafish *tbx5a* wild type (wt) and mutant (*hst* and *oug*) open reading frames were cloned into a pCS2 vector and 300ng of each construct was transfected along with the reporter constructs and normalization control. Experiments were performed in triplo, each with hextuplicate biological replicates. Isolation of cell extracts and subsequent luciferase assays were performed 48h after transfection using Luciferase Assay System according to the protocol of the manufacturer (Promega). Luciferase measurements were performed using a Promega Turner Biosystems Modulus Multimode Reader luminometer. Mean luciferase activity and standard deviation were plotted as fold activation compared to the promoter-reporter plasmid. All data was statistically validated using a one-way ANOVA for all combinations.

### Imaging

In vivo phenotypic assessment and imaging was carried out on a Leica M165FC stereomicroscope or a Zeiss StemiSV6 stereomicroscope (Carl Zeiss AG, Oberkochen, Germany). Embryos were sedated if necessary with 16 mg/ml tricaine (MS222; Sigma-Aldrich) in E3 medium. ISH imaging was performed using a Zeiss Axioplan microscope (Carl Zeiss AG). Images were captured with a DFC420 digital microscope camera (Leica Microsystems). Confocal imaging was carried out on a Leica SPE or SP8 confocal microscope (Leica Microsystems). Multiphoton imaging was carried out on a Leica SP5 or SP8 confocal microscope (Leica Microsystems). Time-lapse imaging was carried out on sedated, PTU-treated, *tnnt2a* morpholino oligo-injected and dechorionated embryos mounted in 0.25% agarose in E3 medium. Images were acquired using a Leica SP5 or SP8 multiphoton microscope and stacks were acquired approximately every 10min for about 16h.

### Image analysis and statistical assays

Time-lapse: Imaris software (Oxford Imaging) was used to generate time-lapse movies and automated cell tracking in 3D, followed by manual inspection of individual tracks. Time lapse movies spanned approximately 28 hpf-42 hpf, with a frame (full stack) acquisition period of approximately 13 min. For each movie analyzed, tracks were selected if they were contained a minimum of 15 acquisition points. Drift correction was applied in Imaris prior to track analysis to correct for displacement of the whole heart during image acquisition.

Calculation of the track displacement angle θ: the axis of the heart was drawn as a line through the middle of the dorsal venous and arterial poles of the heart at the first frame of the time lapse (28 hpf). For each track, θ is the angle formed by the displacement vector of a track with the heart axis as indicated in Fig. 2J. All data presented in the manuscript on time-lapse movies were generated in Imaris and subsequently processed in Excel (Microsoft) if required.

Cell roundness: cell roundness assessment was carried out in Fiji freeware (www.fiji.sc). Roundness of a cell is defined as:

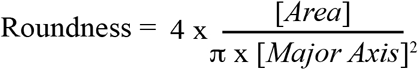

Cell counting: cell counting was carried out in Volocity (Quorum technologies) on confocal-acquired 3D stacks.All statistical assays were carried out in Graphpad Prism 6.0 (GraphPad Software).

## Acknowledgements

The authors thank Anko de Graaff (Hubrecht Imaging Center) for assistance with microscopic imaging and Phong Nguyen (Hubrecht Institute) and Hessel Honkoop (Hubrecht Institute) for critically reading the manuscript.

## Funding

The authors wish to acknowledge the support from the Dutch Heart Foundation grant CVON2014-18CONCOR-GENES to J.B. and V.M..

## Competing Interests

The authors declare no competing or financial interests.

## Author Contributions

Conceptualization: F.T., V.M.C., J.B.; Methodology: F.T., V.M.C., J.B.; Validation: F.T., F.K.; Formal analysis: F.T., F.K., S.C.v.d.B., M.v.d.B; Investigation: F.T., F.K., S.C.v.d.B., M.v.d.B; Writing-original draft: F.T., M.v.d.B., J.B.; Supervision: F.T., V.M.C., J.B.; Funding acquisition: V.M.C., J.B.

## Supplementary Material

**Figure S1.**
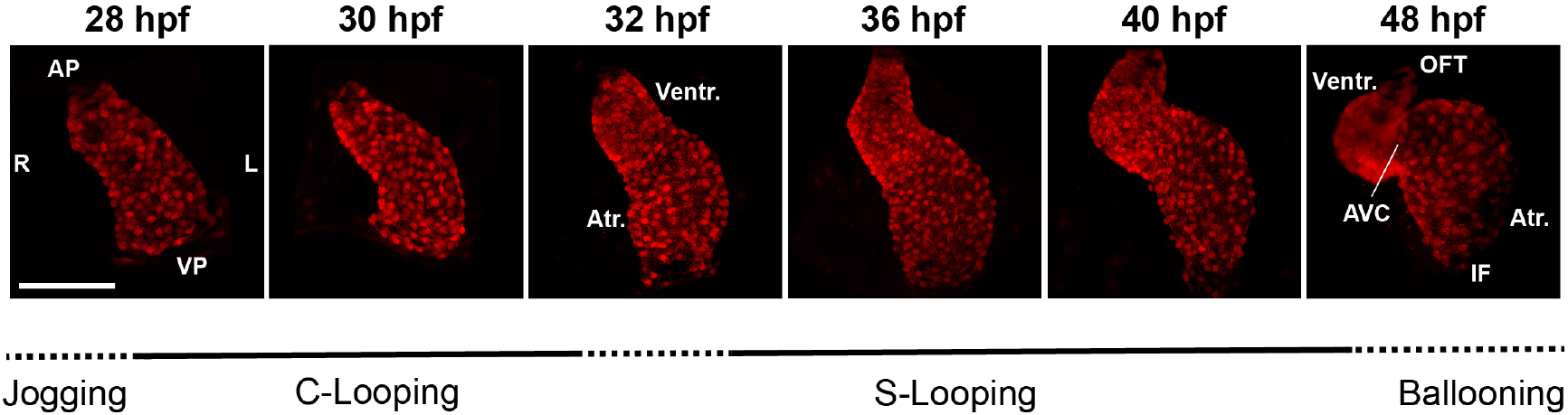
Zebrafish Cardiac Looping. Representative Z-stack projections of *tg(myl7:Gal4FF; UAS:RFP)* hearts at 28 hpf, 30 hpf, 32 hpf, 36 hpf, 40 hpf and 48 hpf. VP: Venous Pole; AP: Arterial Pole; Atr.: Atrium; Ventr.:Ventricle; IF: Inflow; OFT: Outflow Tract; AV canal: Atrio-Ventricular Canal. Scale bar: 100 μm.

**Figure S2.**
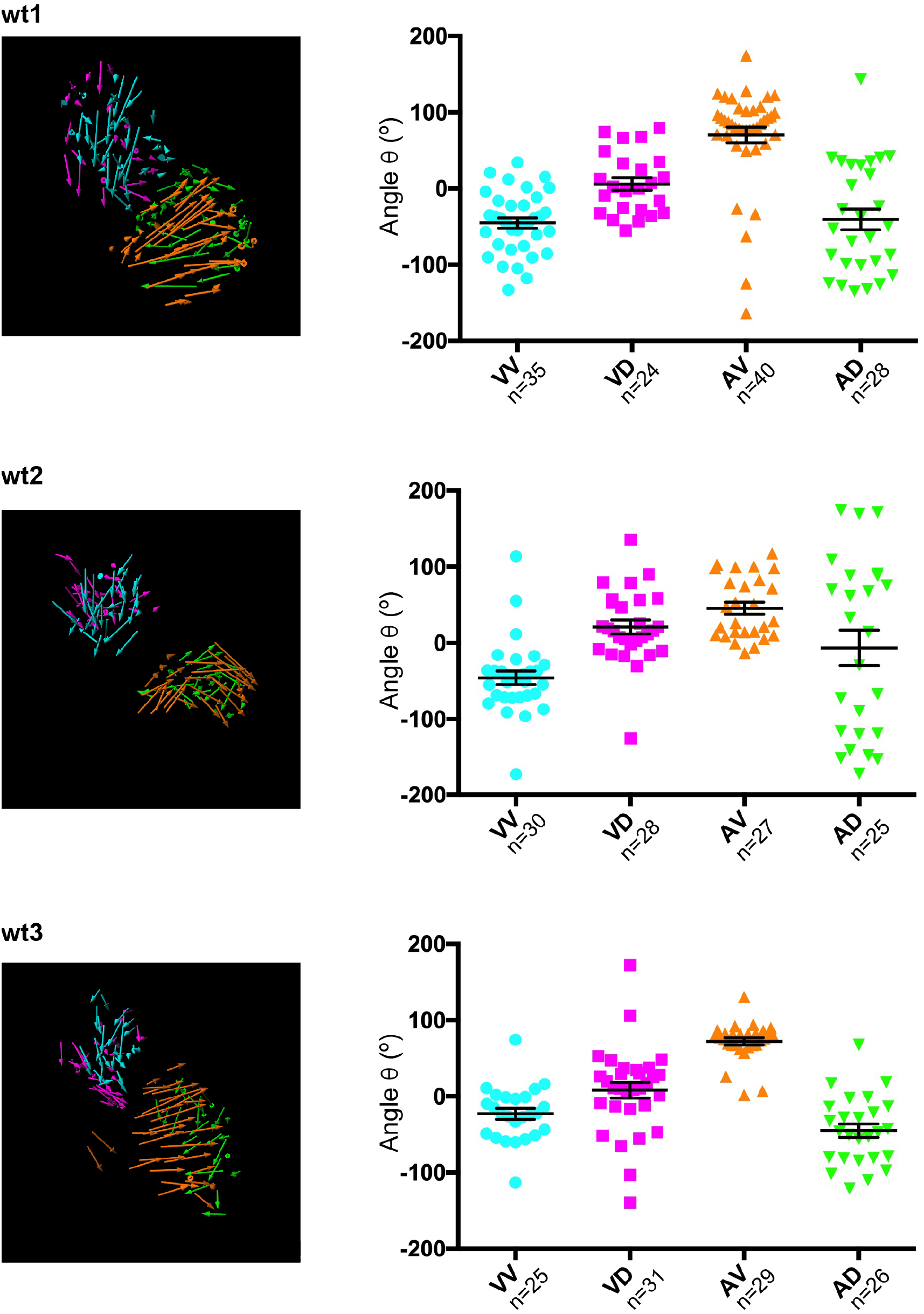
Single data points for angle of displacement measurements in wt hearts. Left panel: Displacement tracks for different heart regions. Right panels: Bars: mean value +/− SEM. VV: Ventricle Ventral; VD: Ventricle Dorsal; AV: Atrium Ventral, AD: Atrium Dorsal.

**Figure S3.**
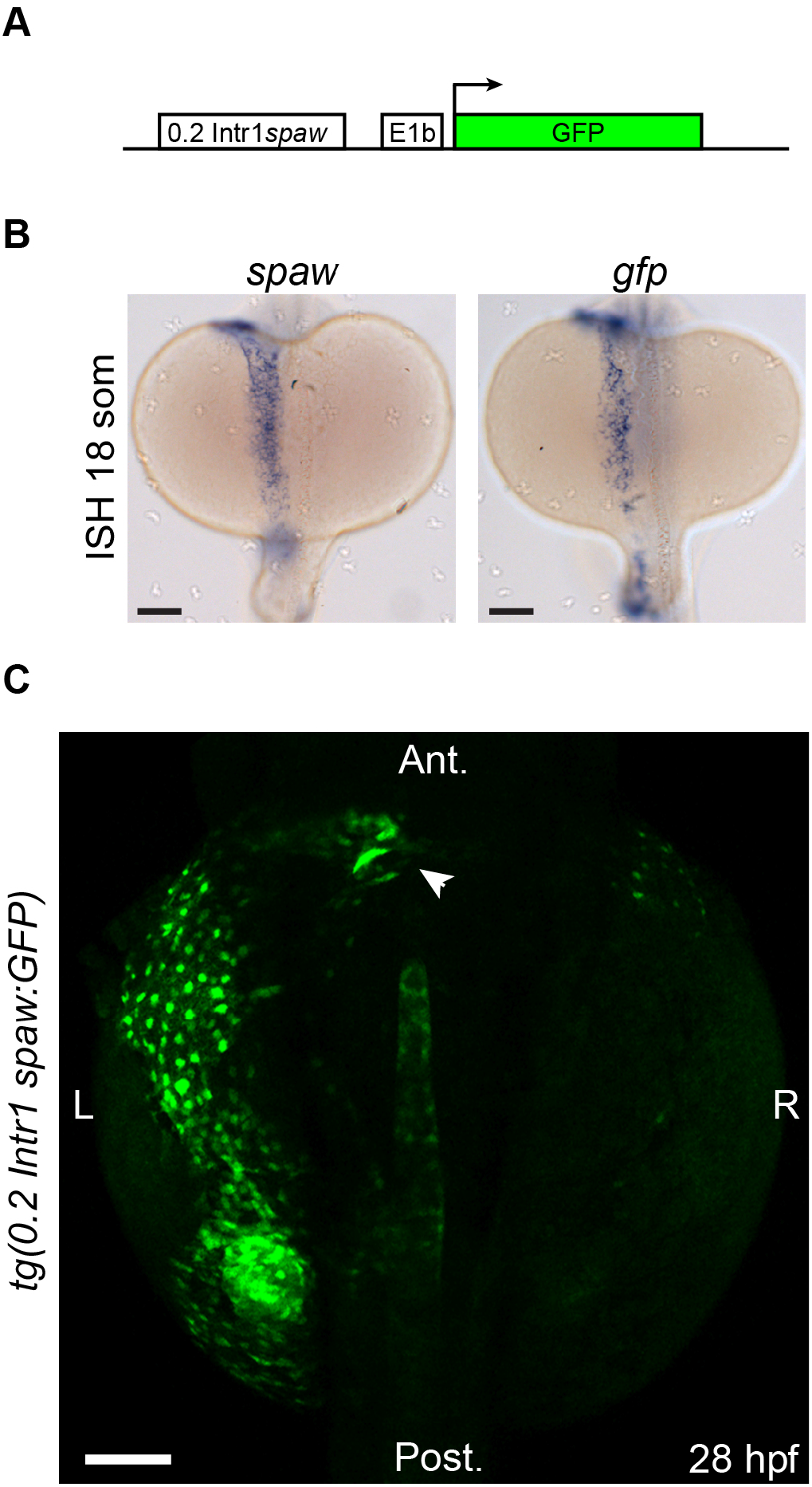
Transgenic *0.2Intr1spaw:eGFP* reporter line. **(A)** An approximately 0.2 kb conserved sequence located in Intron 1 of the *spaw* genomic locus, hereafter called 0.2Intr1spaw, was used to drive expression of GFP. **(B)** ISH on *spaw* and *GFP* at 18 somites confirms the validity of the tg(0.2Intr1spaw:eGFP) reporter expression pattern. **(C)** GFP fluorescence at 28 hpf illustrates the left LPM reporter use of the *0.2Intr1spaw:eGFP* line. Arrowhead indicates arterial pole of the heart tube. Scale bar: 100 μm.

**Figure S4.**
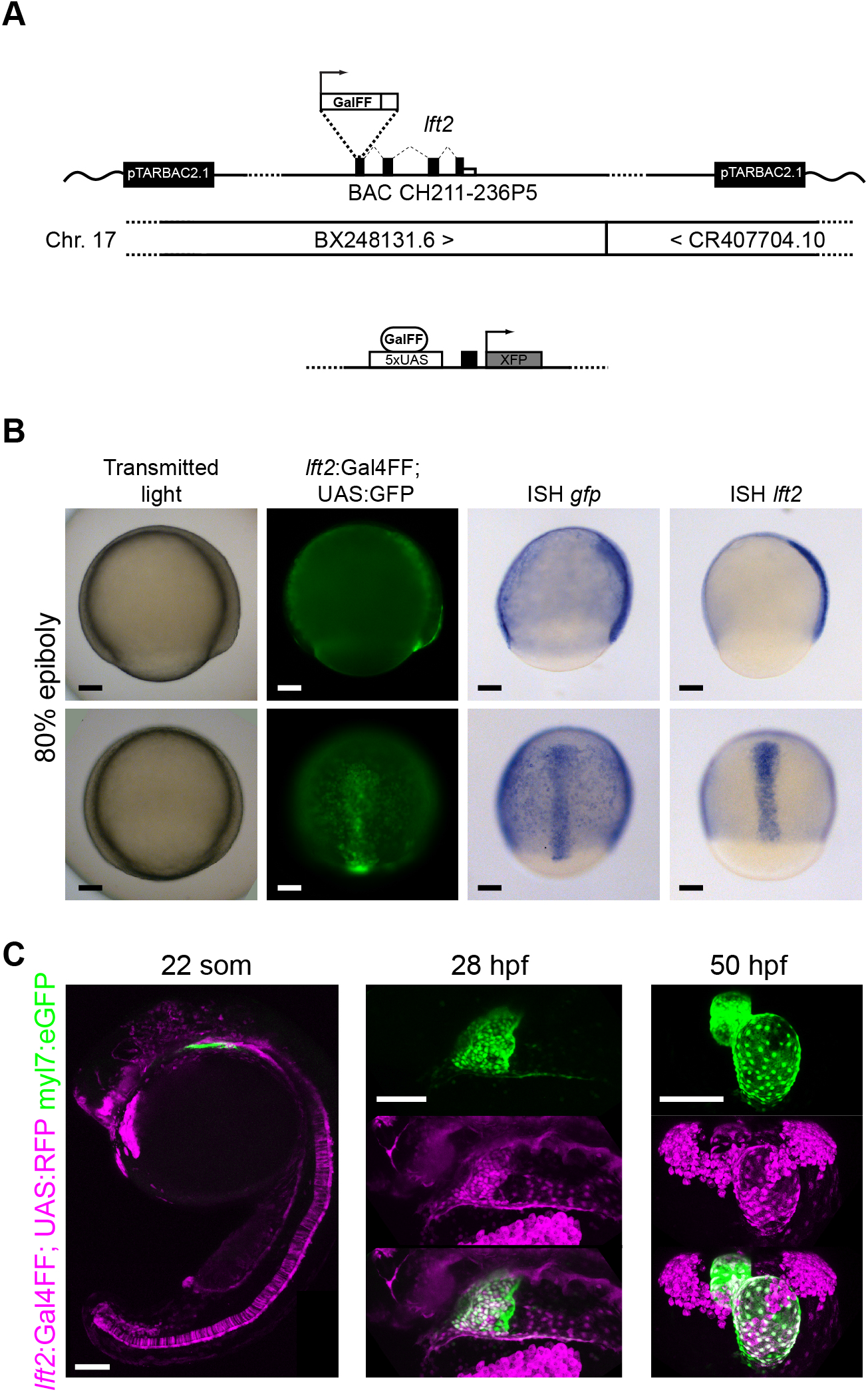
Transgenic *lft2BAC:Gal4FF* reporter line. **(A)** An expression cassette containing Gal4FF was recombined into BAC CH211-236P5 at the ATG site of the 1^st^ exon of the *lft2* gene. **(B)** GFP expression pattern of the *tg(lft2BAC:Gal4FF; UAS:GFP)* line at 80% epiboly and ISH for *GFP* and *lft2*. Note the overlap of expression at the forming midline. **(C)** Reporter expression (RFP) of the *tg(lft2BAC:Gal4FF)* reporter line at in the whole embryo at 22 somites (lateral view, left panel) and in the cardiac region at 28 hpf (center panel, lateral view) and at 50 hpf (right panel, frontal view). Scale bar: 100 μm.

**Figure S5.**
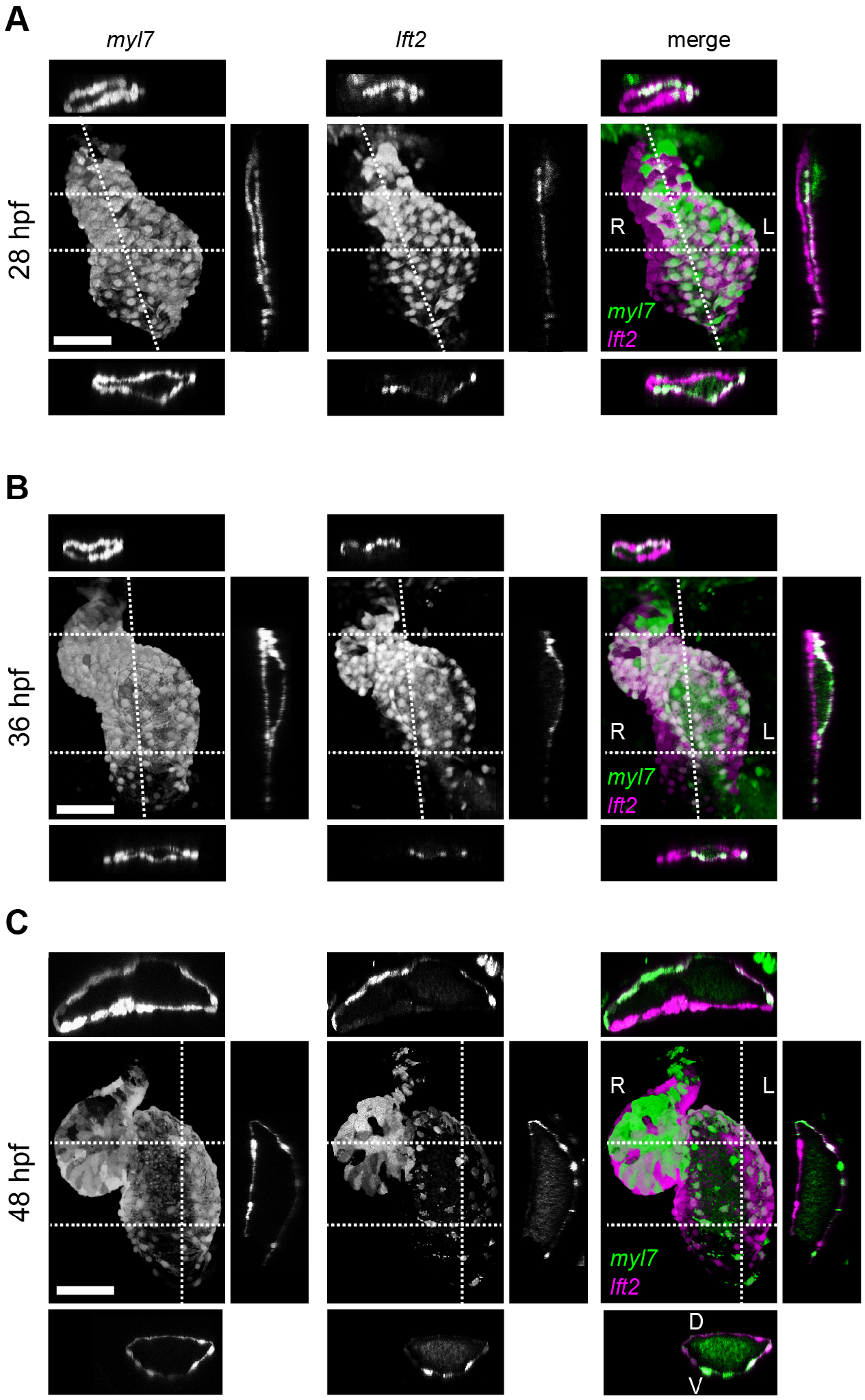
Use of the *lft2BAC:Gal4FF* to track left-and -right originating cardiomyocytes during cardiac looping. Compare panels with Fig.2. **(A)** At 28 hpf, left- and right-originating cardiomyocytes are organized in a ventral, dorsal manner, as reported previously (38). **(B)** As cardiac loping progresses, left-originating cells are displaced towards the outer curvatures of the ventricle and atrium. Conversely, at the inner curvature of the atrium only right-originating cells can be observed. **(C)** After completion of cardiac looping, left-originating cells are located at the outer curvatures of the ventricle and atrium, as also observed in the *0.2Intr1spaw:eGFP* line. Scale bar: 50 μm.

**Figure S6.**
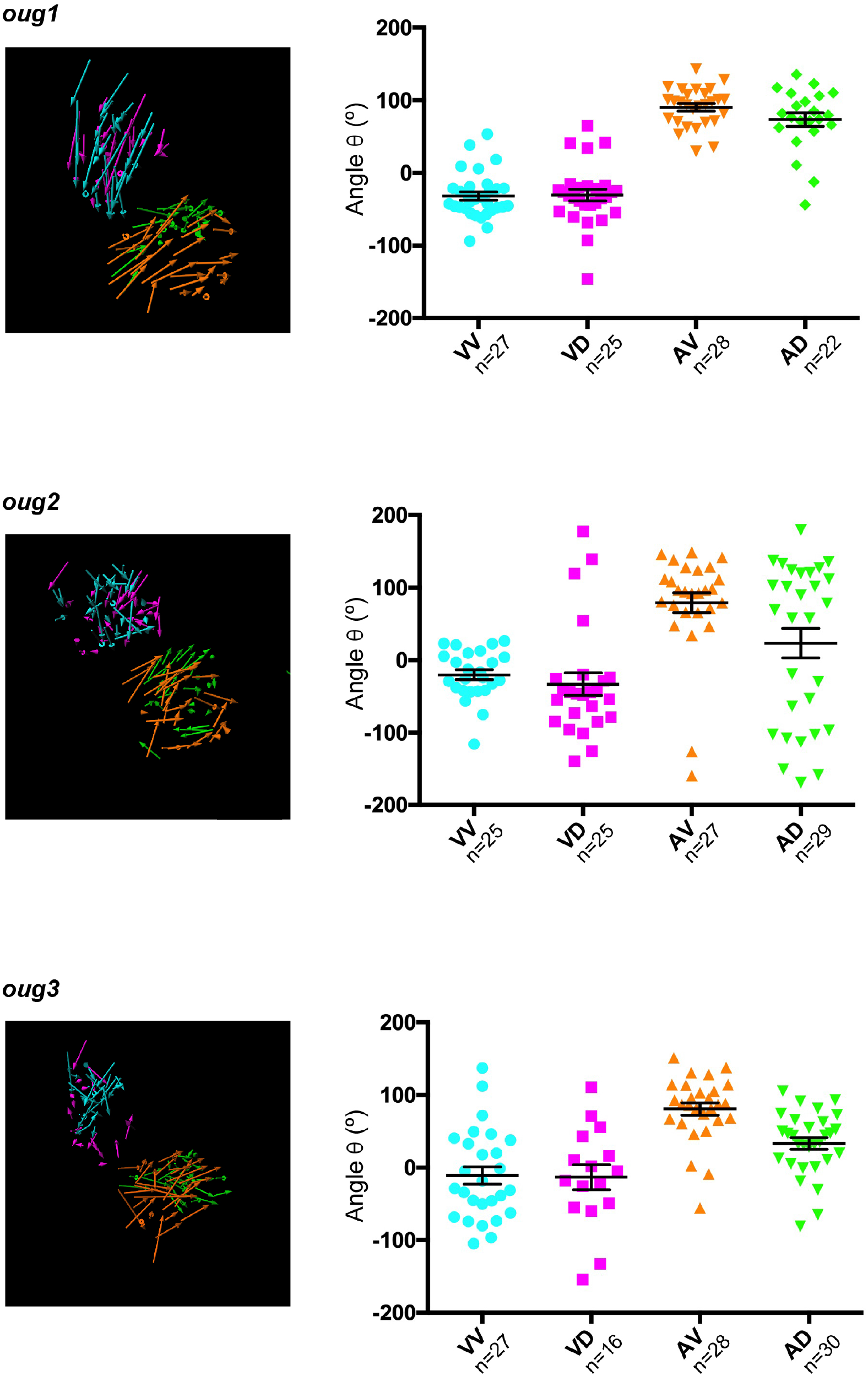
Single data points for angle of displacement measurements in *oug* hearts. Left panel: Displacement tracks for different heart regions. Right panels: horizontal bars: mean value +/− SEM.VV: Ventricle Ventral; VD: Ventricle Dorsal; AV: Atrium Ventral, AD: Atrium Dorsal.

**Figure S7.**
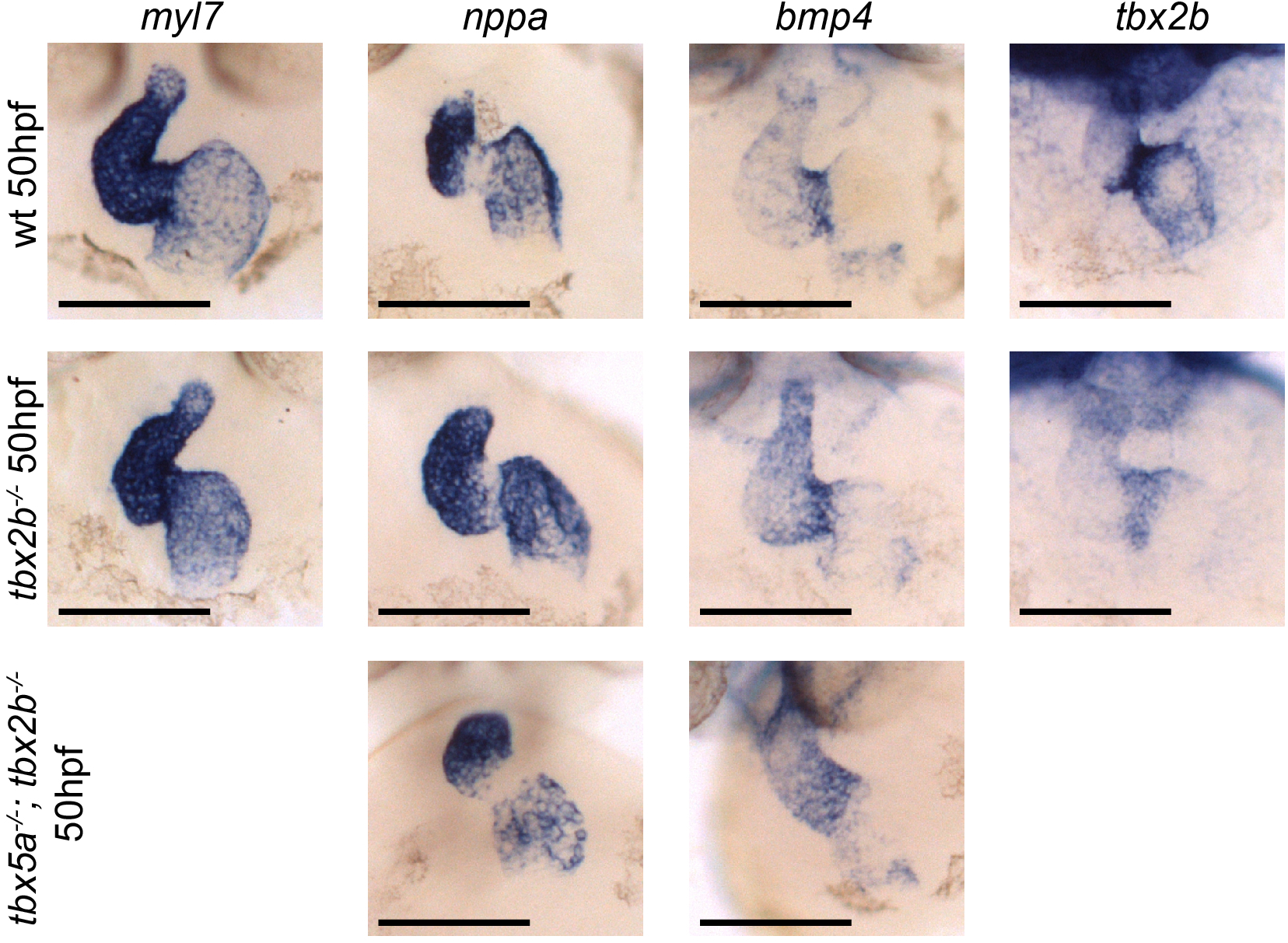
Analysis of cardiac markers in *fby/tbx2b*^*-/-*^ *and tbx5a*^*-/-*^; *tbx2b*^*-/-*^ embryos at 50 hpf. ISH probes used are *myl7* (all cardiomycytes), *nppa* (cardiac chambers), *bmp4* (AV canal and IFT) and *tbx2b* (AV canal). Scale bar: 100 μm.

**Figure S8.**
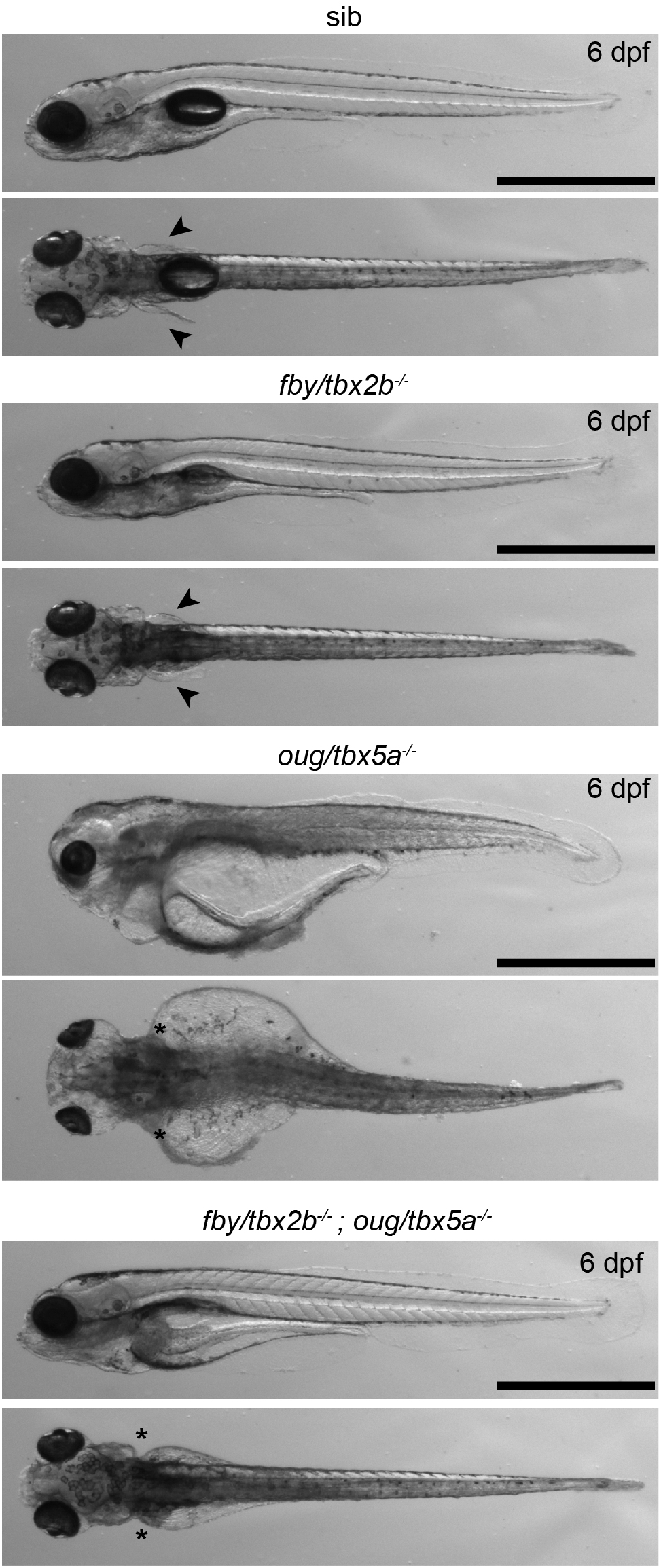
Phenotypical analysis of *tbx5a*^*-/-*^; *tbx2b*^*-/-*^ larvae at 6dpf. Arrowheads indicate presence of pectoral fins in wt (sib) **(A)** and *tbx2b*^*-/-*^ **(B)** larvae, asterisks indicate absence of pectoral fins in *tbx5a*^*-/-*^ **(C)** and *tbx5a*^*-/-*^; *tbx2b*^*-/-*^ **(D)** larvae. Note that the severe general oedemic phenotype displayed by *tbx5a*^*-/-*^ larvae is rescued in the *tbx5a*^*-/-*^; *tbx2b*^*-/-*^; double mutant. Scale bar: 500 μm.

## Notes

### Competing Interest Statement

The authors have declared no competing interest.

